# Exon inclusion signatures enable accurate estimation of splicing factor activity

**DOI:** 10.1101/2024.06.21.600051

**Authors:** Miquel Anglada-Girotto, Carolina Segura-Morales, Daniel F. Moakley, Chaolin Zhang, Samuel Miravet-Verde, Andrea Califano, Luis Serrano

**Affiliations:** Centre for Genomic Regulation (CRG), The Barcelona Institute of Science and Technology, Dr. Aiguader 88, Barcelona 08003, Spain; Department of Systems Biology, Vagelos College of Physicians and Surgeons, Columbia University Irving Medical Center, New York, USA 10032; Department of Biochemistry & Molecular Biophysics, Vagelos College of Physicians and Surgeons, Columbia University Irving Medical Center, New York, USA 10032; Center for Motor Neuron Biology and Disease, Columbia University, New York, USA 10032; Department of Biology, Institute of Microbiology and Swiss Institute of Bioinformatics, ETH Zurich, Zurich, Switzerland; Department of Medicine, Vagelos College of Physicians and Surgeons, Columbia University Irving Medical Center, New York, USA 10032; Herbert Irving Comprehensive Cancer Center, Columbia University Irving Medical Center, New York, USA 10032; Chan Zuckerberg Biohub New York, New York, NY, USA; Department of Biomedical Informatics, Vagelos College of Physicians and Surgeons, Columbia University Irving Medical Center, New York, USA 10032; Universitat Pompeu Fabra (UPF), Barcelona, Spain; ICREA, Pg. Lluís Companys 23, Barcelona 08010, Spain

**Keywords:** cancer, alternative splicing, splicing factor, protein activity, VIPER, carcinogenesis

## Abstract

Splicing factors control exon inclusion in messenger RNAs, shaping transcriptome and proteome diversity. Their catalytic activity is regulated by multiple layers, making single-omic measurements on their own fall short in identifying which splicing factors underlie a phenotype. Here, we posit that splicing factor activity can be estimated from changes in exon inclusion. To test this hypothesis, we benchmarked methods for constructing splicing factor→exon networks and estimating splicing factor activity. We found that combining RNA-seq perturbation-based networks with VIPER (Virtual Inference of Protein Activity by Enriched Regulon analysis) accurately captures splicing factor activation as modulated by multiple regulatory layers. This approach integrates splicing factor regulation into a single score derived solely from exon inclusion signatures, allowing functional interpretation of heterogeneous conditions. As a proof of concept, we identify recurrent cancer splicing programs, revealing oncogenic- and tumor suppressor-like splicing factors missed by conventional methods. These programs correlate with patient survival and key cancer hallmarks: initiation, proliferation, and immune evasion. Altogether, we show splicing factor activity can be accurately estimated from exon inclusion changes, enabling comprehensive analyses of splicing regulation with minimal data requirements.

## INTRODUCTION

Splicing factors carry out alternative splicing: they selectively regulate intron removal from expressed precursor mRNAs. Together with alternative transcription start and termination sites, splicing factors determine each gene’s pool of mature mRNAs, impacting overall mRNA stability, translation, protein activation, and protein-protein interactions^1^. Simultaneously, splicing factor activity is regulated by multiple mechanisms, including transcription of their encoding genes, pre-mRNA splicing, protein synthesis and degradation, post-translational modifications, subcellular localization, and formation of functional complexes with other proteins (Fig. 1). Accurate assessment of splicing factor activity is thus challenging as it requires accounting for highly diverse regulatory mechanisms.

**Figure 1.**
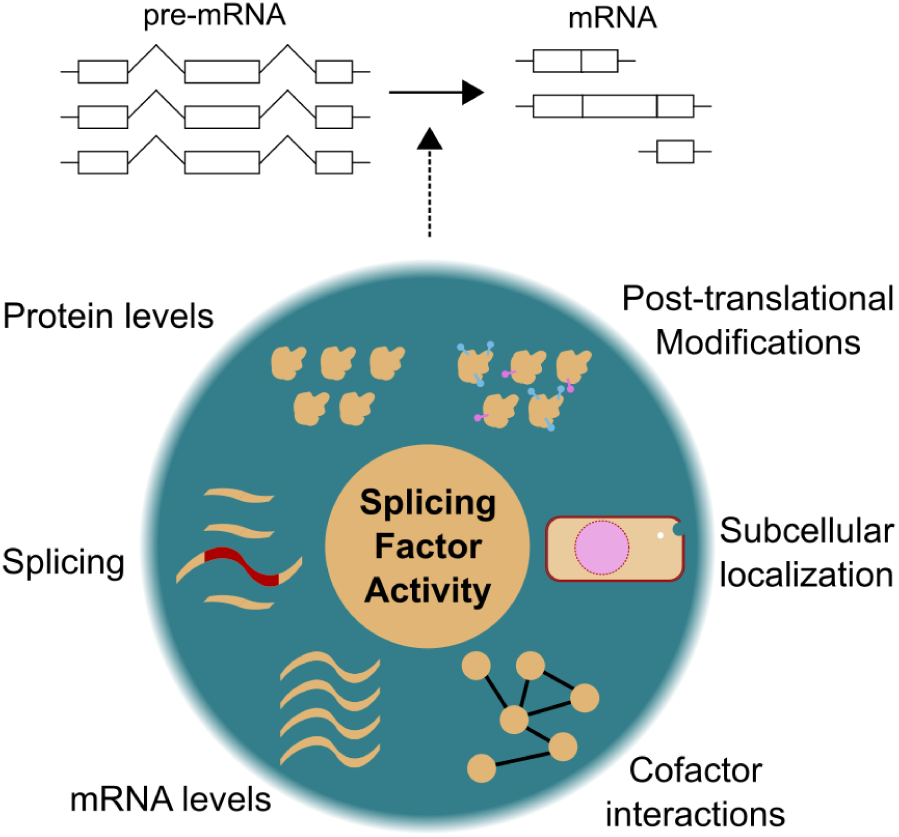
Splicing factor activity is determined by multiple levels of regulation. Splicing factors regulate the removal of introns from pre-mRNA. The activity of splicing factors is regulated by multiple molecular mechanisms such as their mRNA levels, alternative splicing, protein levels, post-translational modifications, subcellular localization, and cofactor interactions.

Splicing factor activities have been typically inferred from the presence of activating and inactivating somatic mutations or differential gene expression^2–5^. However, these represent just two of the many possible mechanisms contributing to splicing factor activity modulation. This partial picture limits identifying aberrant splicing factor activity, especially in complex genetic contexts, such as cancer, where multiple molecular alterations contribute to cancer hallmarks^4,6–9^. A possible solution could be collecting more omic measurements on splicing factors. However, this would not only require large-scale efforts and resources but also methods to integrate these data and calculate splicing factor activities, which are still elusive.

The challenge of assessing protein activity has been effectively addressed for transcription factors^10^. Rather than relying on the omics of a transcription factor to estimate its activity, focusing on the expression of its target genes reflects the complex combined influence of regulatory and signaling mechanisms that determine its activity. This requires addressing two problems: first, assessing the transcriptional targets of every transcription factor, and second, estimating their activity based on the differential expression of these targets. Transcription factor→gene interactions (i.e., regulatory networks) can be inferred computationally from gene expression datasets by reverse engineering algorithms such as ARACNe^11,12^, as well as empirically, by mining curated databases generated from publicly available experiments^13–15^. Then, given a high-quality regulatory network that includes the probability (likelihood) and directionality (mode of regulation) of each interaction, transcription factor activity can be accurately assessed from expression signatures through bioinformatic methods like VIPER^16^ and others^17^. Although this approach only requires transcriptomic data, it has been proven highly valuable in disentangling the common functional states of transcriptional regulatory circuits in heterogeneous molecular contexts like cancer^6^.

Following the same reasoning, we posit that exon inclusion signatures report on the activity of splicing factors. If this is the case frameworks like VIPER could be extended to estimate splicing factor activity from changes in exon inclusion across their target mRNAs, represented as networks of splicing factor→exon interactions (splicing factor networks, for simplicity) (Fig. 2a). To do so we need to identify which approaches for network construction and activity estimation best adapt protein activity methods to splicing factors, and whether this can uncover novel insights from heterogeneous data such as cancer samples, where multiple regulatory layers are simultaneously dysregulated.

**Figure 2.**
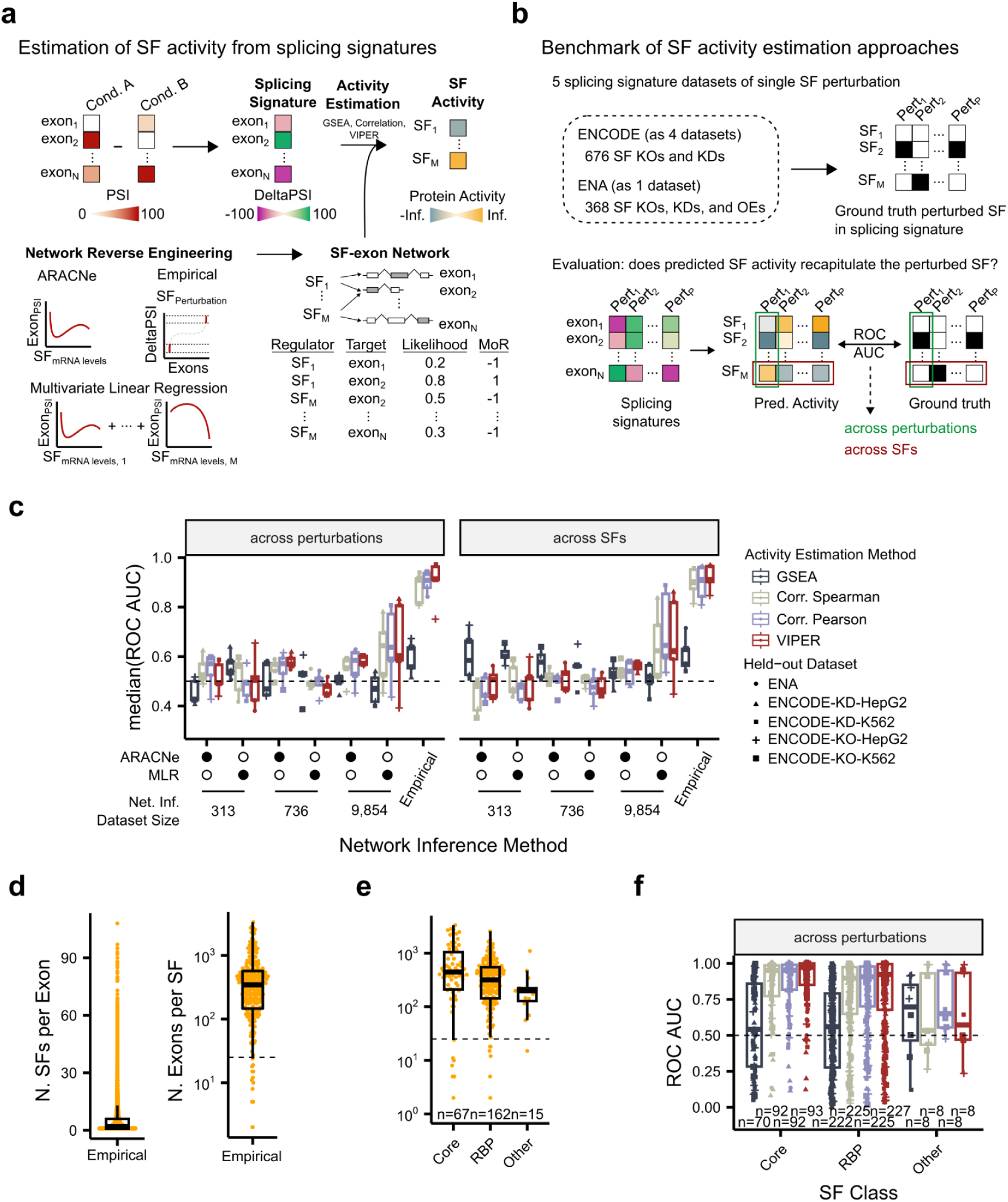
Reverse engineering of splicing factor networks to estimate splicing factor activities with VIPER. (**a**) Estimation of splicing factor activity from exon inclusion signatures. The algorithm requires two inputs: (i) a splicing signature, the difference between exon inclusion in a condition of interest (Cond. A) and a reference condition (Cond. B); (ii) a splicing factor (SF) network, the splicing factor→exon interactions and their corresponding likelihood and mode of regulation (MoR). Splicing factor networks can be obtained using either empirical –from splicing factor experimental perturbations– or computational –here we consider ARACNe and Multivariate Linear Regression (MLR)– methods. We estimated splicing factor activity using normalized enrichment scores (NES) from gene set enrichment analysis (GSEA), Pearson and Spearman correlation coefficients, and VIPER. (**b**) Benchmark outline for splicing factor activity estimation algorithms. We generated 5 benchmark datasets including single-splicing factor perturbation experiments from different publicly available sources. To compute performance metrics (ROC-AUC), we consider predicted splicing factor activities and which splicing factor was actually perturbed across perturbations (horizontal) and across splicing factors (vertical). (**c**) Evaluation of splicing factor activity estimation with different methods for activity estimation from an exon inclusion signature and a splicing factor→exon network. Computational networks (ARACNe, MLR) were reverse-engineered using three different datasets of increasing size. For "Empirical" networks, performance reflects evaluation on a held-out benchmark dataset, with networks constructed using all others. Each benchmark dataset consists of independent experiments, potentially from the same or different cellular contexts relative to the held-out sets. Among these, only when ENA is held out can we assess performance across truly independent cellular contexts for "Empirical" networks. See Supplementary Fig. 2b for this specific evaluation. (**d**) Distribution of number splicing factors per target exon (left), and number of target exons per splicing factor (right) in empirical splicing factor networks. (**e**) Distributions of the number of targets (exons) per regulator (splicing factor) considering either all splicing factors categorized by “Core” splicing factors appear in Spliceosome Database^40^, “RBP” splicing factors that appear in Gene Ontology^78^ category “RNA binding protein”, or “Other” splicing factors. (**f**) Evaluation of splicing factor activity estimation for each splicing factor in benchmark datasets using empirical splicing factor networks. In box and whisker plots in panels (**c**), (**d**), (**e**), and (**f**), the median is marked by a horizontal line, with the first and third quartiles as box edges. Whiskers extend up to 1.5 times the interquartile range, and individual outliers are plotted beyond.

Here, to repurpose protein activity analysis for splicing factors, we benchmark different methods for splicing factor network inference and activity estimation. We find that combining empirically derived splicing factor networks with the VIPER framework yields the most accurate activity estimates across our benchmark datasets. Notably, this approach remains effective even in experiments that are more complex that those used to construct the networks. To illustrate its broader applicability, we applied the framework to cancer-related phenotypes. This uncovered two biologically relevant and complementary splicing programs that are not detected by standard splicing factor expression analysis. Integration with transcriptomic, proteomic and phosphoproteomic data suggests that these cancer-associated splicing programs are coordinately regulated through diverse molecular alterations spanning multiple regulatory layers. Altogether, splicing factor activity analysis is a powerful method for generating testable hypotheses about the splicing factors that drive complex cellular phenotypes.

## RESULTS

### Empirical networks outperform computational ones for activity estimation

To evaluate the accuracy of splicing factor activity estimation methods, we created a set of benchmark datasets based on splicing factor perturbation experiments based on a consensus list of 509 splicing factors compiled from the literature (Supplementary Table 1 and 2)^8,18–21^. We curated 474 bulk RNA-seq experiments perturbing 248 distinct splicing factors and computed exon inclusion changes (“Delta PSI”) relative to control samples. These datasets allowed us to compute ROC-AUC scores to benchmark whether estimated activities correctly identify the perturbed splicing factor from exon inclusion signatures (Supplementary Note 1; Supplementary Fig. 1a-b).

We compared computational and empirical strategies for constructing splicing factor networks, which serve as input to activity estimation. Computational networks were inferred from large RNA-seq datasets using ARACNe and multivariate regression, while empirical networks were constructed from observed exon inclusion (PSI) changes exceeding 15% upon splicing factor perturbation. Specifically, to define empirical networks, we explored a range of thresholds on exon inclusion changes and tested their impact on performance. Although the exon targets in empirical networks may reflect both direct and indirect regulatory effects, they are functionally linked to the splicing factor’s activity.

We evaluated splicing factor activity estimation with VIPER alongside three baselines (gene set enrichment analysis (GSEA), Spearman and Pearson correlations) on both computational and empirical networks. Each method interprets splicing factor networks differently: GSEA uses the network as an unweighted gene set, correlations incorporate interaction likelihood and regulatory direction, and VIPER combines both through a rank-based enrichment analysis (aREA). Across all benchmarks and estimation methods, empirical networks combined with VIPER consistently outperform other strategies (Fig. 2b-c), indicating that experimentally derived networks are more effective for capturing true splicing factor activity.

Full details on network construction, threshold optimization, and benchmarking are provided in Supplementary Note 2 and Supplementary Fig. 1c.

Additionally, we found that the poor performance of computational networks is largely due to their inaccurate prediction of regulatory direction, which limits their utility for activity estimation (Supplementary Fig. 1d-f). Full details of this analysis are available in Supplementary Note 3.

### Splicing factor activity estimation is robust across studies and contexts

To assess the robustness of VIPER-based activity estimation, we evaluated performance across the diverse studies and experimental contexts represented in our benchmark datasets. Despite the well-documented context-specificity of certain splicing changes, we found that empirical networks combined with VIPER consistently achieve high accuracy (ROC-AUC ∼ 0.9), indicating strong generalizability (Supplementary Fig. 2a-b).

To better understand the basis of this robustness (i.e., the stability, reliability, and consistency of the method across diverse conditions), we examined the structure of empirical networks. Each exon is typically regulated by a small number of splicing factors (median = 2), indicating this combination of functional splicing factor→exon interactions is commonly highly specific despite the fact that in some cases the same exon can be regulated by 90 or more splicing factors (Fig. 2d). Indeed, when reducing the number of interactions considered, accuracy remains high, confirming that a reduced subset of interactions is sufficient for reliable activity inference (Supplementary Fig. 2c).

We also assessed how robustness varies by splicing factor class. Core spliceosomal factors, those directly involved in the splicing reaction, tend to be more highly connected (Fig. 2e). Benchmarking by factor class revealed that activity estimates for core splicing factors achieve higher accuracy than those for other types such as RNA-binding proteins (RBPs) (Fig. 2f). This likely reflects that the functional targets of core factors are more stable across conditions, whereas RBP targets are more context-dependent and may therefore vary across datasets, reducing inference accuracy (median ROC-AUC_VIPER_: core = 0.97; RBP = 0.92).

Of note, while most benchmarked splicing factors (i.e., those perturbed in more than one dataset) achieved ROC-AUC values > 0.5, in certain cases the perturbed factor showed AUC values below 0.5 (e.g., Supplementary Fig. 2a). This may indicate that, in those conditions, activities estimated from the held-out dataset are inconsistent with the expected effect. Biologically, this could be due to co-regulation: perturbing one splicing factor may elicit stronger effects on upstream regulators or downstream effectors. Technically, benchmarking across datasets could introduce batch effects or experimental noise. These single-factor perturbation benchmarks therefore need to be complemented by validations using orthogonal perturbation approaches, as we describe below.

As we did not find independent replicates for all perturbed splicing factors, we chose to retain all available factors in our benchmarks, even when quality metrics in this particular benchmark may be suboptimal, to provide a broad, comparative view of method performance, highlight incomplete networks or pathways, and enable hypothesis generation in less-characterized regions of the splicing machinery.

Finally, to further confirm the consistency of our findings, we repeated all benchmarking analyses using an alternative metric: the rank percentile of the perturbed factor among all factors. These rank percentiles showed strong agreement with the ROC-AUC-based results (Extended Data Fig. 1), reinforcing the robustness of our conclusions.

A detailed description of these robustness analyses is provided in Supplementary Note 4.

### Empirical networks capture functional regulation

Splicing factor perturbations can influence exon inclusion indirectly through intermediate regulators. For instance, not all splicing factors bind RNA; yet their perturbation often produces consistent exon inclusion signatures. To investigate the extent and impact of such indirect regulation in empirical networks, we compared them to crosslinking and immunoprecipitation (CLIP)-based networks^22^, which capture direct RNA-binding events. We found that most empirical interactions were not supported by CLIP, indicating a predominance of indirect regulation. Nevertheless, VIPER maintains high predictive accuracy even when restricted to empirical networks containing only indirect interactions (Supplementary Fig. 2d-f).

These results demonstrate that direct binding evidence is not essential for accurate activity estimation, underscoring the robustness of the functional interactions captured by empirical networks. Full details on CLIP network construction and analysis are provided in Supplementary Note 5.

### Estimated splicing factor activity recapitulates regulatory mechanisms of increasing complexity

We have demonstrated above that perturbation-derived splicing factor networks combined with VIPER can identify which splicing factor was perturbed under similar experimental conditions. However, we propose that this approach goes beyond that, offering the ability to functionally capture complex multi-omic regulatory mechanisms that ultimately influence splicing factor activity solely through exon inclusion signature analysis (Fig. 1). To illustrate this, we evaluate splicing factor activity analysis across several experimental validations involving perturbations of diverse regulatory mechanisms.

For example, the drug Indisulam downregulates the protein level of splicing factor RBM39 by mediating its proteasomal degradation (Fig. 3a and b)^23,24^. In contrast, the mRNA levels of its encoding gene increase consistently upon treatment, probably as a compensatory mechanism (Supplementary Fig. 3)^23,24^. In this case, using changes in splicing factor mRNA levels to evaluate splicing factor activity would lead to the incorrect conclusion that Indisulam increases RBM39’s activity. In sharp contrast, VIPER analysis of transcriptional profiles following drug perturbation identifies RBM39 as the splicing factor with the greatest decrease in splicing activity, thus effectively recapitulating Indisulam’s mechanism of action (Fig. 3c). As a result, VIPER-based analysis recapitulated protein-level effects, despite using only transcriptomic data.

**Figure 3.**
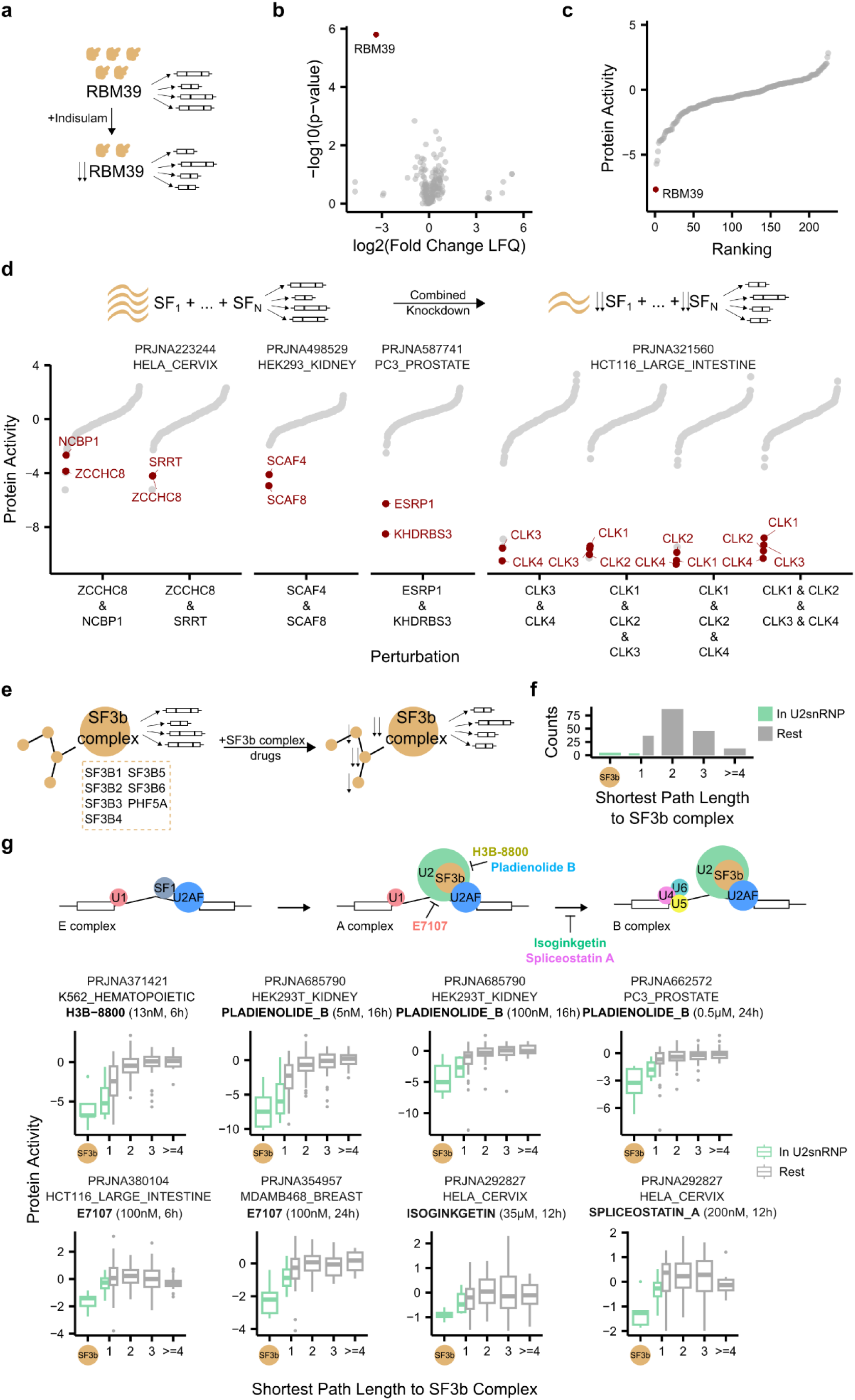
Assessing splicing factor activity in complex molecular contexts. (**a**) Indisulam promotes protein-level degradation of splicing factor RBM39, inducing significant changes in the exon inclusion repertoire of the cell. (**b**) Differential proteomic expression of splicing factors. Relationship between fold change LFQ values comparing Indisulam-treated (5 μM, 6 hours) and untreated samples and p-values obtained from a two-sided T-test in IMR-32 cells. Red, RBM39. (**c**) Ranking of splicing factors according to their estimated protein activities upon treatment with Indisulam (5 μM, 6 hours) in IMR-32 cells. Red, RBM39. (**d**) Distributions of estimated protein activities upon combinatorial perturbation of splicing factors. Red, corresponding knocked down genes in each case. Top, experimental outline. Studies analyzed include PRJNA223244^28^, PRJNA498529^27^, PRJNA587741^26^, and PRJNA321560^25^. (**e**) The SF3b complex is composed of 8 proteins. When perturbing spliceosome assembly with drugs, we hypothesize proteins in the SF3b complex and proteins interacting with the complex should decrease their activity. Adapted from Rogalska *et al.*^8^ (**f**) Distribution of shortest path length to reach any protein of the SF3b complex in the STRINGDB protein-protein interaction network. Color, whether genes belong to the U2snRNP. (**g**) Top, mechanisms of action of splicing-targeting drugs: H3B-8800 and Pladienolide B bind the SF3b complex; E7107 prevents the binding of U2 complex to RNA; Isoginkgetin and Spliceostatin A impede the transition from A to B complex. Bottom, distributions of splicing factor activities upon perturbation with different splicing-targeting drugs considering the shortest path lengths of each splicing factor to the SF3b complex. Color, whether genes belong to the U2snRNP. Studies analyzed include PRJNA371421^30^, PRJNA685790^65^, PRJNA662572^66^, PRJNA380104^67^, PRJNA354957^68^, and PRJNA292827^69^. In box and whiskers plots in the panel, the median is marked by a horizontal line, with the first and third quartiles as box edges. Whiskers extend up to 1.5 times the interquartile range, and individual outliers are plotted beyond.

Splicing factors are often modulated in concert, leading to complex and intermingled splicing signatures. To evaluate whether our approach could recapitulate the effect of such combinatorial perturbations, we used VIPER to analyze the transcriptomic data from four different studies where from two to four splicing factors were simultaneously silenced^25–28^ (Fig. 3d). Confirming the value of the proposed methodology, the silenced splicing factors were always among the most inactivated. This suggests that combinatorial splicing factor perturbations produce additive changes in exon inclusion, thus allowing effective assessment of splicing factor activity by our approach (Fig. 3d).

Splicing factors interact to form a large and dynamic protein complex called the spliceosome^21^. We thus hypothesized that the VIPER-assessed activity of splicing factors interacting in the spliceosome complex via protein-protein interactions (PPIs) should be correlated. At the beginning of the splicing reaction, during splice site recognition, the spliceosome forms the E, A, and B complexes comprising small ribonucleoprotein (snRNP) complexes (U1, U2, U4, U5 and U6)^8^. Within the U2 snRNP, the SF3b complex (SF3B1, SF3B2, SF3B3, SF3B4, SF3B5, SF3B6, PHF5A) plays a pivotal role in branch point recognition^29^. Different splicing drugs perturb this step of the reaction. For instance, H3B-8800 and Pladienolide B bind SF3B1^30,31^; E7107 prevents the binding of U2 snRNP to RNA^32^; and Isoginkgetin and Spliceostatin A prevent the transition from A to B spliceosome complexes in the splicing reaction^33,34^. It is thus reasonable to expect that perturbing the SF3b complex with these drugs should strongly decrease the activity of the associated splicing factors, proportionally to their distance to the SF3b complex in the PPI network (Fig. 3e and f). Consistent with this prediction, treating cell lines with the aforementioned drugs induced notable VIPER-assessed inactivation of splicing factors in the SF3b complex, followed by splicing factors in the U2 snRNP and other splicing factors with direct (shortest path length in PPI space = 1) and indirect (shortest path length in PPI space > 1) interactions with the SF3b complex (Fig. 3g).

Taken together, these data further demonstrate that combining VIPER with empirical splicing factor networks accurately recapitulates splicing factor activity modulation mechanisms across a range of diverse perturbations that were not used to generate the splicing factor networks.

### Recurrent aberrant splicing factor activity defines two cancer splicing programs

Cancer co-opts the splicing machinery in favor of tumor progression^4,9,35^. However, few studies have examined whether these splicing programs are unique to specific cancer cohorts or are shared, with most research focusing on single regulatory layers of splicing factors^36,37^.

Our approach, based solely on exon inclusion changes, captures various regulatory mechanisms driving splicing factor activity. This suggests that estimated splicing factor activities reflect the combined output of all regulatory inputs affecting their function. To demonstrate its utility, we investigated whether splicing factor dysregulation in cancer follows a general pattern or varies by cohort. We analyzed multiple cancer cohorts to determine if certain splicing factors are consistently dysregulated, suggesting a recurrent cancer splicing program.

Of the 33 cancer cohorts included in The Cancer Genome Atlas (TCGA), 14 include ≥ 10 exon-level transcriptional profiles from primary tumor (PT) and solid tissue normal (STN) biopsies. We leveraged these samples to assess splicing factor activity by comparing each PT sample to the median STN profiles and vice versa (Fig. 4a–b). We then focused on the splicing factors that were significantly differentially active (FDR < 0.05, two-sided Wilcoxon rank sum test) in PTs compared with their corresponding STNs across all cancer cohorts (Fig. 4b). Interestingly, some splicing factors emerged as recurrently activated or inactivated across a majority of cancer cohorts (Fig. 4c; Supplementary Fig. 4a). This suggests that oncogenic alterations may cause coordinated and recurrent dysregulation of splicing factor activity, despite their well-described heterogeneity. Considering a threshold of at least 5 out of 14 cancer cohorts to be indicative of recurrent dysregulation (see Methods), the analysis identified a set of (n=61) putatively oncogenic splicing factors (i.e., aberrantly activated in ≥ 5 cohorts), and a set of (n=61) putatively tumor suppressor splicing factors (i.e., aberrantly inactivated in ≥ 5 cohorts) (Supplementary Table 3). Of note, PTBP1, a splicing factor with confirmed oncogenic roles in four different cancer types^38^, emerged as the most frequently activated splicing factor.

**Figure 4.**
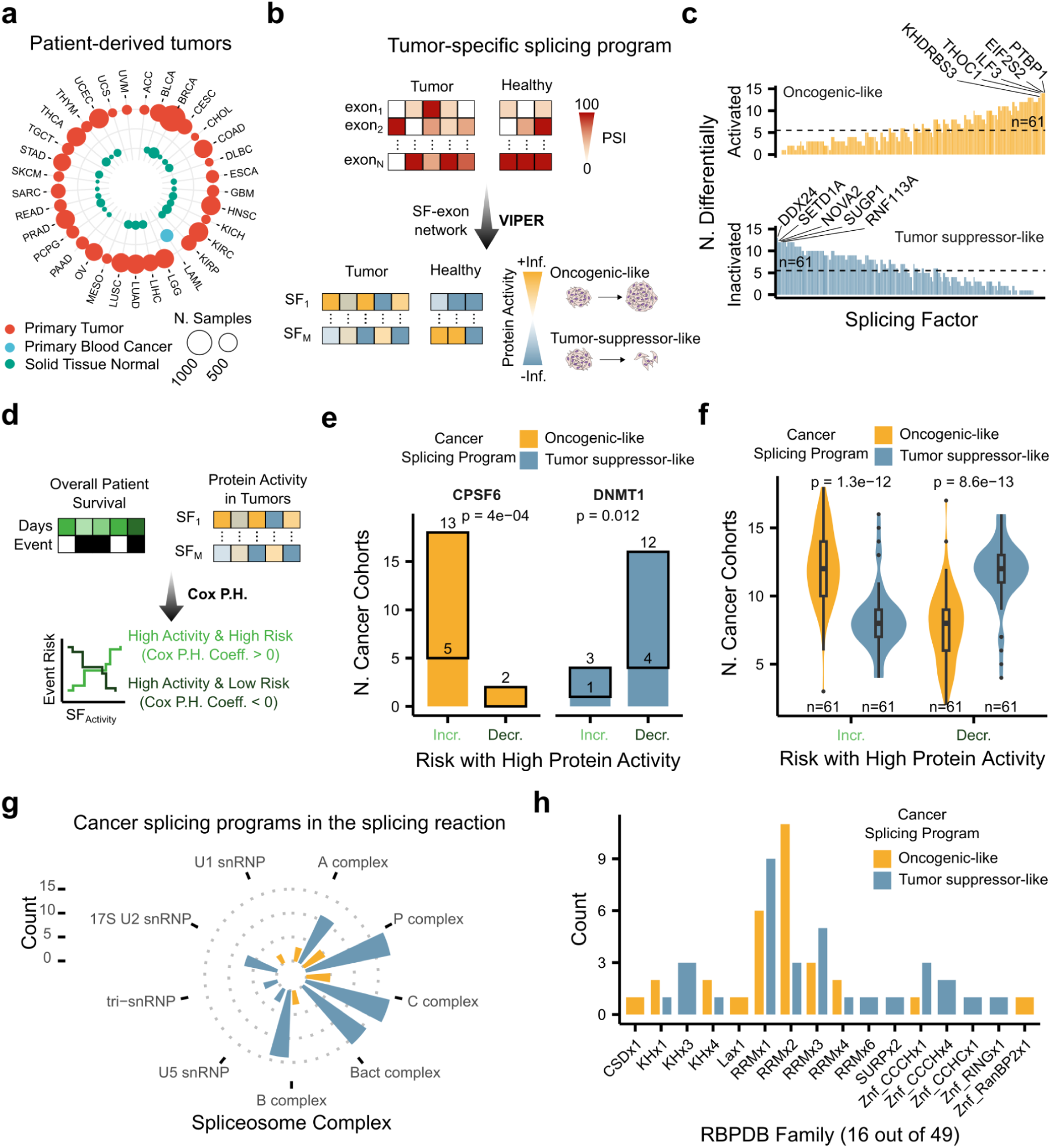
Splicing factors recurrently active in tumors exhibit cancer driver-like behavior. (**a**) TCGA cancer cohorts overview. Among the 33 cohorts with tumor samples, 14 also have corresponding healthy (solid tissue normal) samples. Cancer type abbreviations are available at https://gdc.cancer.gov/resources-tcga-users/tcga-code-tables/tcga-study-abbreviations. (**b**) Workflow to identify tumor-specific splicing programs. From healthy and tumor samples, we quantified their exon inclusion and computed the splicing signatures to obtain splicing activities for each splicing factor and sample. (**c**) Frequency of splicing factors showing significant activation (top) or inactivation (bottom) in tumor samples versus solid tissue normal (FDR < 0.05, two-sided Wilcoxon rank sum tests). Dashed lines mark the recurrence threshold for a splicing factor’s inclusion in either the oncogenic-like or tumor suppressor-like cancer splicing program, with selected splicing factors above each line. Labels, top 5 most recurrently activated or inactivated splicing factors. (**d**) Workflow to associate protein activities in primary tumor samples with patient overall survival using a Cox Proportional Hazards (Cox P.H.) survival model. Positive Cox P.H. coefficients indicate that high splicing factor activity is associated with a high risk of death. Conversely, negative Cox P.H. coefficients indicate that high splicing factor activity is associated with a low risk of death. (**e**) Number of cancer cohorts in which either CPSF6 (left) or DNMT1 (right) are associated with either high or low risk of death. Top, number of cancer cohorts for each type of association. Thick border, number of cancer cohorts with Cox Proportional Hazards coefficient significantly different than 0 (FDR < 0.05). (**f**) Distributions of the number of cancer cohorts that each splicing factor is associated with a high or low risk of death. Top, p-value based on two-sided Wilcoxon rank sum tests. Bottom, number of splicing factors in the corresponding cancer splicing program. (**g** and **h**) Splicing factors from cancer splicing programs identified found in (**g**) the spliceosome reaction complexes according to SpliceosomeDB^40^, and in (**h**) RNA-binding protein (RBP) domain families. In box and whiskers plots in panel (**f**), the median is marked by a horizontal line, with the first and third quartiles as box edges. Whiskers extend up to 1.5 times the interquartile range, and individual outliers are plotted beyond.

If these recurrently dysregulated splicing factors play a role in tumorigenesis and progression, then activation of oncogenic and tumor suppressor factors should be predictive of poor and good prognosis, respectively. Indeed, outcome analyses confirmed these expectations (Fig. 4d; see Methods). For example, analysis of primary tumor samples across 20 TCGA cohorts with sufficient metadata (see Methods) revealed that aberrant activity of a recurrent oncogenic splicing factor (CPSF6) correlates with poor prognosis in 18 cancer cohorts (Cox Proportional Hazards (PH) coefficient > 0) while the activity of a recurrent tumor suppressor splicing factor (DNMT1) correlates with favorable outcome in 16 cancer cohorts (Cox PH coefficient < 0) (Fig. 4e).

Extending the analysis to all 122 recurrently dysregulated splicing factors showed the same trends. Indeed, aberrant activation of oncogenic splicing factors correlates with poor outcomes across significantly more cancer cohorts than tumor suppressor ones (*p* = 1.3·10-^12^, two-sided Wilcoxon rank sum test). The opposite happens for tumor suppressor-like splicing factors; their activity correlates with good outcome across significantly more cancer cohorts than the oncogenic ones (*p* = 8.6·10-^13^, two-sided Wilcoxon rank sum test) (Fig. 4f). These patterns are conserved if we consider different thresholds to define cancer splicing program recurrence (Supplementary Fig. 4b).

Typically, splicing factor activity is extrapolated from expression. We argue that splicing factor expression is only one of the molecular mechanisms regulating splicing factor activity. Hence, we expect that our approach should capture activity differences that are not necessarily reflected by differences in mRNA levels. To evaluate this, we repeated the same analysis with splicing factor gene expression. Confirming our expectations, gene expression fold-changes in tumor vs matched normal samples did not correlate with their differential activity (Spearman correlation = −0.047, *p* = 0.042) (Supplementary Fig. 4c). Candidate oncogenic (n = 106) and tumor suppressor splicing factors (n = 30) identified by gene expression analysis (Supplementary Fig. 4d) were not predictive of poor and good outcome (*p* > 0.05 in both cases) (Supplementary Fig. 4e and f). Thus, by focusing on changes in splicing factor targets rather than in splicing factor expression, our approach systematically elucidated the potential contribution of their dysregulation to tumorigenesis and outcome.

To further characterize these cancer splicing programs, we examined their putative roles in the splicing reaction and in RBP families using SpliceosomeDB^39,40^ and RBPDB^39^, respectively. Of the 122 splicing factors in cancer splicing factor programs, 32 and 54 splicing factors were mapped to each database, respectively. Most were part of the C complex in the splicing reaction –involved in the catalytic steps–, and belonged to the RRMx1 and RRMx2 families of RBP domains (Fig. 4g and h).

### Linking cancer splicing program dysregulation to cancer hallmarks

Given the functional relevance of the cancer splicing programs identified through our approach, we wondered which biological processes they may potentially regulate (Fig. 5a). We performed an overrepresentation analysis (ORA) of the genes corresponding to the exons targeted by either oncogenic-like or tumor suppressor-like splicing factors.

**Figure 5.**
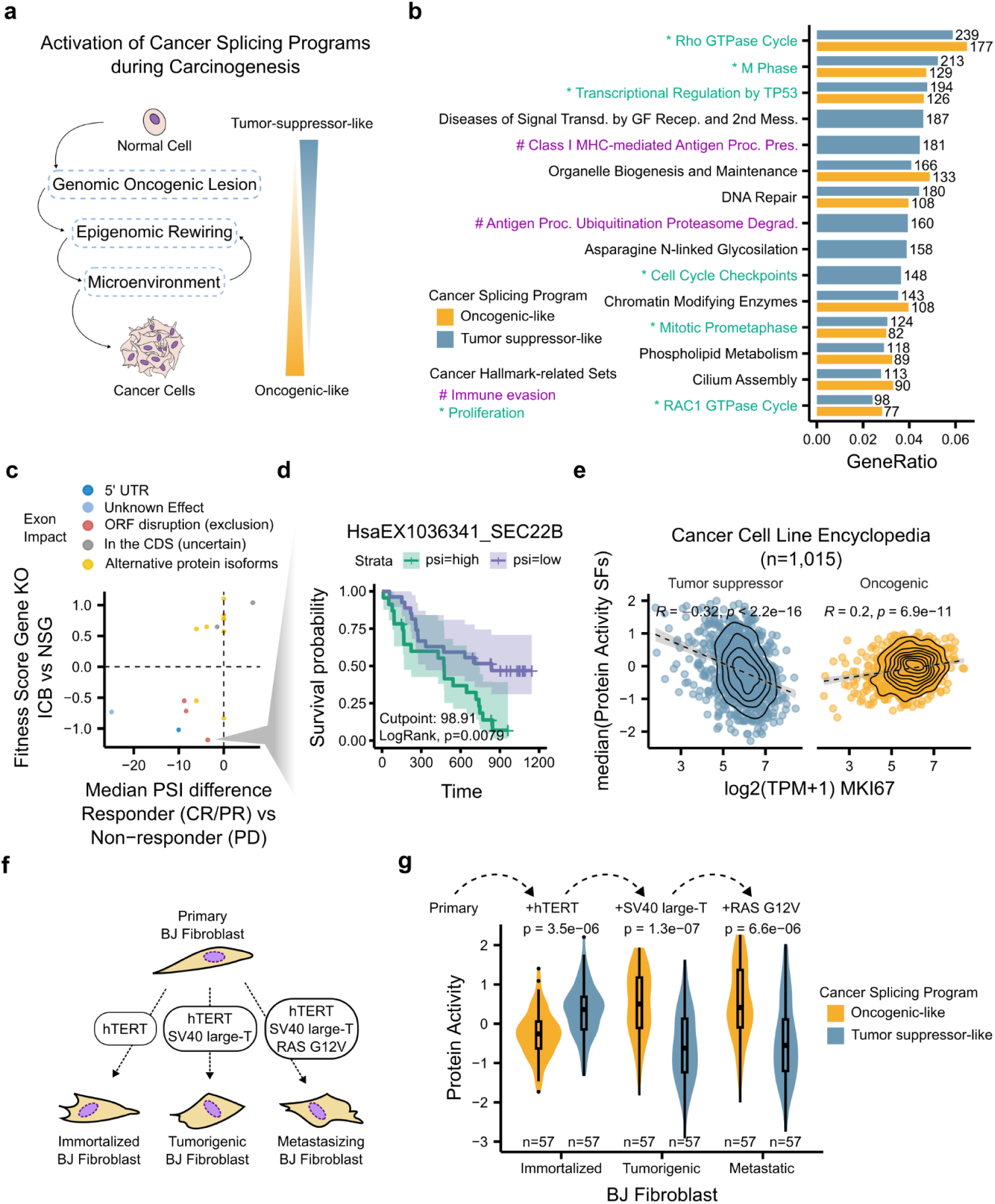
Contribution of cancer splicing programs to disease progression. (**a**) Cancer splicing program activation during carcinogenesis. Carcinogenic transformation begins with genomic lesions that activate oncogenes and inactivate tumor suppressors, reprogramming the cell into a cancerous state. We hypothesize that cancer splicing programs are regulated alongside these transformations. (**b**) ORA of the top 10 significantly (FDR < 0.05) enriched ReactomeDB terms for genes corresponding to target exons of splicing factors in the oncogenic-like and tumor suppressor-like cancer splicing programs. Enriched terms only in the top 10 for one of the splicing programs are also shown. (**c**) Prioritization of exons with the potential to contribute to ICB treatment response. For each of the differentially spliced exons between responder and non-responder patients (*p*-value < 0.05, two-sided Wilcoxon rank sum test), we considered the orthologous gene-level immune evasion score obtained from CRISPR screens comparing the effects in cancer cell viability of knocking out genes in either immunocompetent mice treated with ICB versus immunodeficient mice. Color, exon impact obtained from VastDB annotations. (**d**) Kaplan Meier survival curves of ICB patients in Riaz *et al.*^43^ stratified according to the HsaEX1036341_SEC22B inclusion levels that maximized log-rank test differences. Color, patients with high and low exon inclusion. Bottom, exon inclusion PSI cut point between the two groups of patients and log-rank test *p*-value. (**e**) Relationship between median activities of cancer splicing programs with cancer cell proliferative state. Top, Pearson correlations (R) and corresponding p-values (p) between the variables in the axes. Error shadows, 95% confidence intervals. (**f**) Danielsson *et al.*^46^ experimental design. Primary fibroblasts were induced into different states transfecting either hTERT (immortalized), hTERT and SV40 large-T (tumorigenic), hTERT, SV40 large-T, and RAS G12V (metastasizing). (**g**) Splicing factor activities by cancer splicing program in different stages of carcinogenesis. Top, p-value based on two-sided Wilcoxon rank sum tests comparing the activity of the splicing factors in each cancer program. In box and whiskers plots in panel (**g**), the median is marked by a horizontal line, with the first and third quartiles as box edges. Whiskers extend up to 1.5 times the interquartile range, and individual outliers are plotted beyond.

To investigate the potential involvement of cancer splicing programs in cancer hallmarks, we performed an over-representation analysis (ORA) on genes corresponding to exons targeted by either oncogenic-like or tumor suppressor-like splicing factors. We selected the top 10 enriched terms (FDR < 0.05) with the most target genes for each program, highlighting both shared and unique terms, while ensuring that all significantly enriched terms from each program were included (Fig. 5b; Supplementary Table 4). Two of the enriched terms were related to antigen presentation, a critically regulated process in tumor immunology, and six others were related to cell cycle.

Immune evasion hampers the effectiveness of immune checkpoint blockade (ICB) therapies in cancer patients, complicating treatment response predictions^41^. A total of 181 genes belonging to the immune evasion terms “Antigen Processing Ubiquitination Proteasome Degradation” and “Class I MHC-mediated Antigen Processing Presentation” contain exons regulated by the tumor suppressor-like splicing program (Fig. 5b). To prioritize tumor suppressor-associated exons potentially involved in ICB efficacy, we analyzed exon inclusion differences between ICB-responding and non-responding patients based on RECIST criteria^42^, using data from Riaz *et al.*^43^. Out of 608 exons from genes in immune-related terms, 16 were differentially spliced (*p* < 0.05, two-sided Wilcoxon rank sum tests). For additional prioritization, we used data from an in vivo genome-wide CRISPR screen comparing ICB response tumors in immunocompetent versus immuno-deficient NOD SCID Il2rg−/− control mice^43^, identifying exons within genes affecting ICB response (Fig. 5c). Specifically, exon HsaEX1036341 in SEC22B –one of the strongest immune evasion inhibitors identified (impairment score = −1.2)– emerged as a promising candidate. Exclusion of this exon disrupts SEC22B’s open reading frame, potentially reducing functional protein and improving ICB response, as evidenced by its lower inclusion in responder (PSI = 95.9, n = 10) than in non-responder (PSI = 99.4, n = 22) patients (*p* = 0.024, two-sided Wilcoxon rank sum tests). Survival analysis further supported this finding, with low exon inclusion in ICB-treated patients correlating with better prognosis (*p* = 0.0079, Log-rank test; best cutpoint = 98.91 PSI) (Fig. 5d). Examining empirical networks, we identified three splicing factors –PRPF8, SRSF1, and SF3B3– as regulators of HsaEX1036341. An ORA of their targets revealed enrichment (FDR < 0.05) in cell cycle control pathways and “Class I MHC-mediated Antigen Processing Presentation” (Supplementary Fig. 5a), suggesting that inactivating these splicing factors should impair immune responses as well. In vivo CRISPR data revealed that silencing the mouse ortholog of SF3B3 closely mirrors SEC22B silencing (impairment score = −0.89) compared to silencing SRSF1 and PRPF8 (impairment scores = −0.61 and 0.68, respectively). This suggests SF3B3 as the most likely regulator of HsaEX1036341 in SEC22B. We conclude that analyzing the interactions within the identified cancer splicing programs could unveil exons with potentially relevant roles in ICB efficacy.

To validate the involvement of cancer splicing programs in cell proliferation, we estimated splicing factor activities for 1,015 cell lines in the Cancer Cell Line Encyclopedia (CCLE) and correlated the median activities among either oncogenic-like or tumor suppressor-like splicing factors against MKI67 expression, a marker of cell proliferation^44^. The activity of the oncogenic-like splicing program correlates positively with the proliferation marker (Pearson correlation = 0.2, *p* < 6.9·10-^11^) and the activity of the tumor suppressor-like splicing program correlates negatively (Pearson correlation = −0.32, *p* < 2.2·10-^16^) (Fig. 5e). Further, the sign of these relationships is reproducible when computing the correlation by grouping CCLE cell lines by cancer type of origin (Supplementary Fig. 5b). This not only confirms the link between cell proliferative state and the activation of cancer splicing programs but also corroborates the cancer driver-like behavior of the splicing programs identified in an independent dataset as high proliferation marker expression is simultaneously associated with activation of the oncogenic program and inactivation of the tumor suppressor program.

Taken together, we confirmed the biological relevance of the cancer splicing programs identified by splicing factor analyses exploring their associations with two hallmarks of cancer: immune evasion and aberrant proliferation.

### Cancer splicing programs exhibit coordinated regulation during carcinogenesis

Carcinogenesis is the process through which a normal cell becomes a cancer cell with the potential to form a tumor. This process is typically initiated by the acquisition of cancer-driver mutations that permanently activate oncogenes and inactivate tumor suppressor genes^45^. To achieve full transformation into cancer cells, carcinogenic mutations rewire normal cells at all regulatory levels, including splicing. Then, if cancer splicing programs exhibit a cancer driver-like behavior, they should be modulated to mediate the establishment of the cancerous state during carcinogenesis (Fig. 5a).

To test this, we analyzed how splicing factor activities change during carcinogenic transformation. Danielsson *et al.*^46^ measured the transcriptomes of a model of carcinogenesis consisting of primary BJ fibroblasts immortalized with telomerase reverse transcriptase (TERT), transformed into cancer cells by adding the SV40 large-T antigen, and made metastatic by introducing the oncogenic H-Ras mutation RASG12V (Fig. 5f)^47^. We then estimated the splicing factor activities for the immortalized, tumorigenic, and metastatic cells using primary fibroblasts as the baseline reference (see Methods). While oncogenic-like splicing factors are less active than tumor suppressor-like splicing factors in immortalized cells, oncogenic programs activate while tumor suppressor programs inactivate in transformed and metastatic cells (Fig. 5g).

To identify which molecular changes might drive these coordinated shifts in splicing factor activity, we extended our analysis to include multiple layers of omic measurements on splicing factors. Specifically, we examined changes in splicing factor mRNA expression and splicing (from RNA-seq^46^), protein abundance, and phosphorylation levels (from this study). We found that both transcript and protein levels of splicing factors generally increase with tumorigenesis, independent of their classification as oncogenic or suppressor-like (Supplementary Fig. 6a). Phosphorylation dynamics also intensified, with more splicing factors exhibiting changes in phosphorylation state from the tumorigenic stage (Supplementary Fig. 6b). At the level of splicing, we observed widespread inclusion changes in splicing factors themselves, though no single splicing event or factor type dominated (Supplementary Fig. 6a and b).

To identify which of these multi-omic features best explain changes in splicing factor activity, we fitted regularized multivariate linear models. Using Lasso (strong regularization) and Elastic Net (moderate regularization) regression, we associated 7,575 candidate features, including mRNA levels, splicing event usage, protein abundance, and phosphorylation, with changes in splicing factor program activity (see Methods). Lasso selected 11 predictive features, while Elastic Net identified 98 (Supplementary Fig. 6c), narrowing down the hypothesis space to a manageable number of testable candidates for future studies. These included features such as changes in FUS mRNA levels, protein abundance of RBM23 and SNRPB, phosphorylation sites in YJU2B and RBM23, and splicing alterations in DDX39A (HsaEX0018925 exon) and HNRNPD (HsaINT0077001 intron). These results support that changes in cancer splicing factor activity during carcinogenesis are driven by a multifactorial and coordinated rewiring of multiple regulatory layers. This underscores how a unified functional readout, like splicing factor activity, helps interpreting and prioritizing relevant regulatory changes. Our framework provides such a readout, enabling the generation of mechanistic hypotheses for experimental validation.

Given the regulation of cancer splicing programs during carcinogenesis, we evaluated their potential role as carcinogenic drivers by examining DEMETER2 gene dependency scores from DepMap^48^. Dependency scores summarize gene silencing fitness effects and indicate the cancer driver potential of non-essential genes. Silencing essential genes hampers cell viability broadly, while perturbing non-^49–51^essential tumor suppressors is expected to increase viability (dependency > 0), and silencing non-essential oncogenes should decrease it (dependency < 0)^49–51^. As expected, essential splicing factors have negative dependencies across all cell types and splicing programs (cancerous (n = 626), engineered (HEK, OELE, PRECLH), or fibroblasts (HS729)) (Supplementary Fig. 5c, top). Non-essential, tumor suppressor-like splicing factors exhibit higher dependencies than oncogenic-like ones in fibroblasts (*p* = 0.028, two-sided Wilcoxon rank sum tests) (Supplementary Fig. 5c, bottom). Cancerous and engineered cells show similar trends, but dependencies did not significantly differ between cancer programs (*p* > 0.05, two-sided Wilcoxon rank sum tests). Regulating splicing factors in cancer programs can influence cell proliferation –a cancer hallmark– suggesting that their coordinated regulation could contribute to cancer initiation and progression.

## DISCUSSION

Repurposing VIPER with empirically derived splicing factor networks accurately estimated splicing factor activity solely from inclusion signatures of functional target exons. Although empirically derived splicing factor networks were obtained from independent single-splicing factor knockout, knockdown, or overexpression experiments, our approach recapitulated multi-omic regulation of splicing factors, including protein-level degradation, combinatorial silencing, and SF3b complex protein-protein interactions. This benchmarked exon inclusion signatures as a more universal functional reporter assay of splicing factor activity compared to transcription factors^16^, without having to silence each splicing factor in every possible cellular context to obtain highly predictive VIPER-compatible networks. This predictive power under diverse types of direct and indirect perturbations suggests that we can infer splicing factor activities in conditions as complex as tumor samples. Performing splicing factor activity analysis on primary tumors compared with solid tissue normal samples revealed two new cancer splicing programs with splicing factors that behave like oncogenes and tumor suppressor genes. Our analyses suggest these cancer splicing programs are likely involved in carcinogenesis and disease progression.

Traditional methods to infer aberrant splicing factor activity focus on molecular events affecting their activity, such as the presence of genetic alterations or their differential expression. These not only require multi-omic profiling but also correct assessment of whether each specific event (e.g., a nonsynonymous somatic mutation) may affect splicing factor function. In contrast, splicing factor activity analysis shifts the focus from the splicing factors to the exons they functionally regulate. Although splicing factor networks contain direct and indirect interactions, these are coherent enough for precise assessment of splicing factor activity across experiments of increasing complexity that capture multiple layers of regulation using only transcriptomic data. For example, inducing RBM39 degradation with Indisulam increased RBM39 mRNA levels but decreased its protein levels. While traditional methods would produce conflicting results depending on the omic information available, such as an increase in RBM39 activity based on its increase in mRNA levels or an activity decrease when inspecting its protein levels, the splicing signature of the perturbation assessed through VIPER with empirical splicing factor networks showed a clear decrease in RBM39 activity. Further, only the cancer splicing programs identified through splicing factor activity inference showed disease-relevant behaviors. This places splicing factor activity analysis from exon inclusion signatures as a general approach capable of extracting new and functionally relevant insights from widely available transcriptomic data.

Cancer is a genomically complex disease leading to aberrant cell growth^52^. In cancer cells, the splicing machinery is hijacked, resulting in gene isoforms that initiate and sustain the disease^4,8^. Thousands of changes in splicing occur in cancer cells compared with healthy counterparts, suggesting there is a change in the active programs of splicing factors. While certain splicing factors have shown the ability to initiate cancer (e.g., SF3B1^53^, RBFOX2^54^, PTBP1^55^), the existence of general cancer-driver splicing programs remains underexplored^36^. As a case study, our systematic analysis of cancer samples identified two recurrent splicing factor programs with oncogenic- and tumor suppressor-like behaviors. These programs not only hold prognostic power but also are informative of multiple cancer hallmarks: (i) analysis of splicing factor interactions in our networks highlighted relevant exons and regulators potentially influencing ICB treatment efficacy; (ii) their activation correlates with cell proliferative state; and (iii) they are selectively activated during carcinogenesis. To better understand the regulatory mechanisms underlying carcinogenic changes in activity, we integrated transcriptomic, proteomic, phosphoproteomic, and alternative splicing data of splicing factors. Using regularized multivariate models, we identified candidate features as potential drivers of the splicing factor activity switch. Notably, FUS, classified as oncogenic-like by our framework, ranks highly among predictors, and, although not well-studied in cancer, has demonstrated neurotoxicity when overexpressed^56^, warranting further investigation of its putative role in carcinogenesis. These results not only corroborate the power of splicing factor activity analysis to distill biologically relevant splicing programs in complex molecular contexts but also set the stage for future research on how the activation of these cancer driver-like splicing programs is mechanistically regulated.

A current limitation of our approach to obtain accurate activity estimations is the requirement for splicing factor networks obtained from experiments perturbing single splicing factors. Differently from transcription factors, we showed that the splicing factor networks obtained with ARACNe –the algorithm extensively used to infer transcription factor-gene networks^11,12^– generate informative splicing factor-exon likelihood scores but fail at inferring their mode of regulation sign using bulk RNA sequencing observational data. Interestingly, in a study developed concurrently using the same ARACNe-based pipeline for splicing factor network reconstruction with a single-cell RNA sequencing dataset of mouse cortex, the mode of regulation inference is consistently accurate^57^. This suggests that cell type abundance in bulk RNA sequencing samples could confound the splicing factor-exon associations inferred computationally, and sets the stage for future improvements such as employing single-cell experiments. Nevertheless, we show that splicing factor activities can be accurately inferred for 243 out of the 509 annotated splicing factors using empirical splicing factor networks. These experimentally derived networks can be easily extended as more perturbation experiments become available.

Given the capability of our approach to capture altered splicing factor activity across diverse contexts using only transcriptomic data, we anticipate that splicing factor activity analysis will be instrumental in identifying and studying functionally relevant splicing programs underlying complex phenotypes.

## METHODS

### Compilation of the consensus list of splicing factors

As illustrated in Supplementary Fig. 1a, we combined the splicing factors from 5 different publications^8,18–21^– to create a consensus list of 509 splicing factors. Hegele *et al.*^21^ performed a protein interaction screen among the proteins that copurify with spliceosomes. Papasaikas *et al.*^18^ perturbed many of the components of the splicing machinery. Seiler *et al.*^19^ studied the splicing factors with putative driver mutations across TCGA cancers. Rogalska *et al.*^8^ reviewed the regulation of alternative splicing. Head *et al.*^20^ studied the expression of splicing factors in cancer. This list can be found in Supplementary Table 1.

### Quantification of gene expression and alternative splicing with *vast-tools*

All RNA sequencing samples considered in this study were processed with *vast-tools*^58^ to quantify gene expression and exon inclusion. For each sample, we aligned sequencing reads to the hg38 genome assembly (Hs2 in VastDB) with the command “*vast-tools align --sp Hs2 --EEJ_counts --expr*”. We combined them using “*vast-tools combine --sp Hs2 --keep_raw_reads --keep_raw_incl --TPM -C*”. And, we set to NA the PSI of exons whose inclusion or exclusion was detected with less than 10 reads by running “*vast-tools tidy -min_N 1 -min_SD 0 --min_ALT_use 25 --noVLOW*”. For each dataset, this workflow results in two gene expression tables corresponding to raw transcript counts and transcripts per million (TPM) –these values can range from 0 to potentially infinite– and an exon inclusion table quantifying the percentage of transcripts that include a certain exon or percentage spliced in (PSI) –these values can range from 0 to 100–. The details of each command can be found at https://github.com/vastgroup/vast-tools.

### Assembly of splicing factor perturbation dataset from publicly-available RNA sequencing data

#### ENCORE

The ENCODE subproject called ENCORE^59^ contains baseline and perturbed RNA sequencing data of K562 and HepG2 cell lines targeting RNA binding proteins either through shRNA knockdown or CRISPR knockout. Out of the different RNA binding proteins targeted in ENCORE, 164 overlapped with our consensus list of splicing factors. We then used *vast-tools* to analyze the transcriptomes of baseline and perturbed conditions for the splicing factors.

#### European Nucleotide Archive (ENA)

ENA is a public database of sequencing experiments. To find more splicing factors, we performed an advanced search querying for experimental data with the following filters:

- human samples: *tax_eq(9606)*
- RNA sequencing files: *library_strategy=”RNA-Seq”* and *library_source=”TRANSCRIPTOMIC”*
- title (*sample_title* and *experiment_title*), run (*run_alias*), or sample alias (*sample_alias*) containing any of the splicing factors in the consensus list

This search resulted in a table with 20,659 entries that we hand-curated to only contain experiments that specifically perturbed at least one splicing factor and the corresponding baseline condition for each perturbation. This resulted in 1,597 different experiments measuring the perturbation effects in the transcriptome for 114 splicing factors through knockout, knockdown, and overexpression. See Supplementary Table 2 for more details on each experiment and corresponding ENA identifiers.

### Inference of splicing factor networks

#### Observational datasets for computational inference of splicing factor networks

In this study, we use different algorithms to estimate splicing factor activity from transcriptomic data (see below). These algorithms require a splicing factor network that includes both the likelihood (a score between 0 and 1) and the mode of regulation (positive or negative) of each splicing factor→exon interaction. While splicing factor networks can be derived empirically, they can also be inferred computationally from observational datasets. Here, we generated computationally inferred networks by linking splicing factor gene expression levels with exon inclusion profiles across a large sample set, which provides sufficient variation in splicing factor expression and exon inclusion.

To obtain diverse observational data, we used RNA sequencing datasets from two studies:

- Cardoso-Moreira *et al.*^60^: this dataset includes 313 human samples across 6 tissues (brain, heart, ovary, kidney, testis, and liver) at various stages of development and age.
- The Cancer Genome Atlas (TCGA): this dataset includes RNA sequencing data from 9,854 primary tumor samples and 736 adjacent solid tissue normal samples, offering a wide range of tissue types and cancer cohorts.

We processed these datasets using the *vast-tools* pipeline^58^ to quantify gene expression (as TPM) and exon inclusion (as PSI) as described in previous sections. For each exon, *vast-tools* assigns missing values (NA) when it detects fewer than 10 reads, resulting in some missing data in exon inclusion profiles. Because computational network inference methods cannot operate with missing values, we performed missing data imputation using k-nearest neighbors (KNN) algorithm^61^.

We applied the KNN imputation using the *sklearn.impute.KNNImputer* function with k=5 nearest neighbors, selected to balance data availability with imputation robustness. The imputation estimates the missing PSIs by calculating the Euclidean distance based on non-missing values across samples, filling in each missing PSI using the k closest samples as reference points. Using a moderate k=5 provides robust estimates without over-smoothing the inclusion levels (Supplementary Fig. 7). These imputed PSI values were used exclusively for the computational inference of splicing factor networks.

#### ARACNe

To infer splicing factor networks computationally, we adapted the ARACNe-AP tool^12^. Originally, ARACNe calculates pairwise mutual information (MI) scores between genes in a single matrix. For our purposes, we modified ARACNe-AP to compute MI between two matrices with different row names –one for splicing factor log-normalized gene expression TPMs and another for imputed exon inclusion PSIs. Minimal updates were made to the ARACNe-AP code, and these changes are documented in the repository’s version history (https://github.com/chaolinzhanglab/ARACNe-AP).

We ran ARACNe adapted to two different matrices on each of the observational datasets using default mutual information threshold of 1·10-^8^ and 100 bootstrap iterations. For each bootstrap, we treated the splicing factor expression matrix as the regulator and the exon inclusion matrix as the target, applying data processing inequality specifically to splicing factors to limit their indirect associations. Each bootstrap run generated a network of splicing factor→exon interactions, where MI scores indicated the strength of each interaction.

To refine bootstrapped networks, we kept the top 1,000 exons per splicing factor, ranked by MI score. This selection showed minimal impact on performance across different numbers of top target exons retained (Supplementary Fig. 6). We then consolidated these bootstrap networks using ARACNe’s module with the default p-value threshold of 0.05. This consolidation produced a consensus network containing significant splicing factor→exon interactions with MI values as interaction likelihoods.

To complete the splicing factor networks, we assigned a mode of regulation (values between −1 and 1) to each interaction, indicating the regulatory direction. Following VIPER’s guidelines (https://www.bioconductor.org/packages/release/bioc/vignettes/viper/inst/doc/viper.pdf), we used the *viper::aracne2regulon* function to calculate these values based on the consolidated network’s MI scores. This step created networks with both interaction likelihoods and mode of regulation (inhibition/activation), making them fully compatible with the methods tested for splicing factor activity estimation.

#### Multivariate linear regression (MLR)

To identify pairwise interactions between splicing factors and exons while controlling for other potential splicing factor influences, we applied MLR to each observational dataset. For each exon, we modeled its PSI as the dependent variable, with the standardized, log-normalized TPMs of 509 splicing factors as independent variables. This setup allowed us to determine the contribution of each splicing factor to exon inclusion, producing a regression coefficient and associated p-value for each splicing factor→exon pair.

In these models, the regression coefficient represents the strength of the putative interaction, while the p-value assesses the significance of the interaction. To ensure compatibility with methods used for splicing factor activity estimation, we included only interactions with a nominal p-value below 0.01. For each significant interaction, we used the absolute value of the coefficient as the interaction likelihood and the sign of the coefficient as the mode of regulation (−1 or 1).

This approach produced splicing factor networks with both likelihood and mode of regulation for each splicing factor→exon interaction.

#### Empirical

We constructed empirical splicing factor networks using experimental perturbation data from publicly available sources collected to generate our benchmark datasets. For each perturbation experiment (e.g., splicing factor knockdown, knockout, or over expression), we calculated a delta PSI by subtracting the exon inclusion in the control condition from the perturbed condition. To maintain consistency across perturbation types, we multiplied the delta PSI values by −1 for overexpression experiments, assuming that exons more included upon splicing factor upregulation would be excluded upon downregulation in the same cell system. This approach only affects the sign of the delta PSI values, not the original PSI values, which remain within the 0-100% range.

Next, we selected exons showing substantial inclusion changes (|delta PSI| ≥ threshold) upon splicing factor perturbation. The absolute delta PSI served as the interaction likelihood, while the delta PSI sign indicated the interaction’s mode of regulation. Across thresholds from 5% to 45% (in 5% increments), we assessed the resulting networks’ accuracy by calculating ROC-AUC evaluation metrics. A 15% threshold achieved optimal performance and practical network sizes, encompassing 243 of the 248 splicing factors with available experimental data, with high ROC-AUC scores for VIPER-based activity estimation (Supplementary Fig. 1c).

For splicing factors with multiple perturbation experiments available, we aggregated individual splicing factor networks at the dataset level, preserving independent biological replicates. In the ENA dataset, with multiple independent exon inclusion signatures for some splicing factors, we grouped networks by sets of studies to reflect different experiments.

To ensure comparability with the ENCODE-derived networks –which is separated by cell line (HepG2 or K562) and perturbation type (CRISPR knockdown or shRNA knockdown)– we also categorized ENA-derived networks into four distinct groups of experiments perturbing different splicing factors. This resulted in a total of eight networks:

I. ENCORE knockdowns in HepG2 (41,477 splicing factor-exon interactions, 104 splicing factors);
II. ENCORE knockdowns in K562 (45,396 splicing factor-exon interactions, 104 splicing factors);
III. ENCORE knockouts in HepG2 (6,340 splicing factor-exon interactions, 44 splicing factors);
IV. ENCORE knockouts in K562 (17,119 splicing factor-exon interactions, 61 splicing factors);
V. ENA perturbations in different cell systems group 1 (14,158 splicing factor-exon interactions, 108 splicing factors, 48 studies);
VI. ENA perturbations in different cell systems group 2 (3,042 splicing factor-exon interactions, 23 splicing factors, 9 studies);
VII. ENA perturbations in different cell systems group 3 (522 splicing factor-exon interactions, 12 splicing factors, 3 studies); and
VIII. ENA perturbations in different cell systems group 4 (963 splicing factor-exon interactions, 2 splicing factors, 2 studies)

#### Cross-linking and immunoprecipitation (CLIP)

To construct CLIP-based splicing factor networks, we obtained CLIP peaks from the POSTAR3 database^22^. We associated each splicing factor with exons by mapping CLIP peaks to the nearest exon in VastDB within a 500 bp window. In this network, we assigned both the likelihood and mode of regulation to 1, as CLIP data only confirms the presence of an interaction but does not indicate its interaction probability or regulation direction.

#### ARACNe likelihood and Empirical mode of regulation

To assess whether the likelihood values generated by ARACNe contribute to the low performance in splicing factor activity estimation, we created splicing factor networks by replacing empirical likelihoods with those calculated by ARACNe. We included only splicing factor→exon interactions present in both the empirical and ARACNe-derived networks, ensuring a shared interaction basis for comparison.

#### MLR likelihood and Empirical mode of regulation

Using a similar approach to that applied to ARACNe-derived networks, we substituted the empirical likelihoods with likelihoods derived from MLR. As with ARACNe, we retained only splicing factor→exon interactions common in both empirical and MLR networks to allow a direct comparison.

### Estimation of splicing factor activities using splicing factor networks

We estimated splicing factor activities using four complementary methods: normalized enrichment scores (NES) from gene set enrichment analysis (GSEA), Spearman correlation, Pearson correlation, and VIPER. Each method uses different principles for estimating activity, allowing us to benchmark different approaches for splicing factor activity estimation from a splicing factor network and an exon inclusion signature.

#### NES from GSEA

Using GSEA, we estimated splicing factor activities as NES values. Here, each splicing factor network was used as an ontology where splicing factors represent exon sets and exon signatures are ranked by their inclusion levels. Missing PSIs were omitted. We applied the *clusterProfiler::GSEA*^62^ function with the default *fgsea* algorithm^63^. Likelihood and mode of regulation data could not be included in this algorithm. The GSEA function was set with *pvalueCutoff=1.1* to ensure NES scores were produced for all splicing factors, using default settings for other parameters.

#### Correlation-based activity estimation (Spearman and Pearson correlation)

To capture direct relationships between splicing factors and exon inclusion, we estimated splicing factor activities as correlation coefficients. For Spearman and Pearson correlations, we used *stats::cor*, specifying “spearman” and “pearson” methods, respectively. To incorporate the interaction likelihoods and modes of regulation, each splicing factor network was preprocessed by multiplying these values, producing a refined interaction metric. Correlation of this metric with exon inclusion levels provided activity estimates for each splicing factor.

### VIPER

VIPER predicts splicing factor activity by assessing the enrichment of target exons in splicing signatures. It requires two inputs: a molecular signature, such as differences in exon inclusion between perturbed and unperturbed conditions, and a splicing factor network that specifies each factor’s target exons, along with a likelihood score and a mode of regulation.

To calculate protein activity, VIPER applies the analytic Rank-based Enrichment Analysis (aREA)^16^, testing the enrichment of a regulator’s target exons in the input molecular signature. If any exon inclusion values are missing, VIPER omits these values during the enrichment analysis.

VIPER evaluates the significance of activity predictions by comparing enrichment scores to a null model generated through samples permutation. When sample size is limited, VIPER uses permutation or an analytic approximation to assess enrichment. It provides activity scores for all network regulators as output.

### Evaluation metrics of splicing factor activity estimation

To evaluate the accuracy of different methods for estimating splicing factor activity, we constructed benchmark datasets from publicly available single-splicing factor perturbation experiments. These datasets included all relevant experiments from ENCODE and ENA that perturb a single splicing factor, each producing an exon inclusion signature tied to a specific splicing factor’s alteration. We divided these experiments into five benchmark datasets based on source ENCODE-KD-HepG2, ENCODE-KD-K562, ENCODE-KO-HepG2, ENCODE-KO-K562, and ENA.

The goal is for a reliable network and estimation method to accurately identify which splicing factor was perturbed in each experiment. For example, given an exon inclusion signature from a knockdown experiment, the best-performing approach should correctly assign the lowest activity score to the knocked-down splicing factor.

To evaluate performance, we calculated area under the receiver operator curve (ROC-AUC) scores using the *pROC::roc* function^64^, comparing performance across two perspectives: by splicing factor and by perturbation (Fig. 2b). First, we estimated splicing factor activities for each perturbation signature, resulting in a table where rows represent splicing factors and columns represent perturbation experiments.

To calculate ROC-AUC scores across splicing factors, we used each row in this table as a predictor (the activity scores) and a binary indicator marking whether the splicing factor was perturbed as the response. This tests how well the method identifies a perturbation for each splicing factor across all experiments. Similarly, we calculated ROC-AUC scores across perturbations by using each column as a predictor and a binary vector marking the perturbed factor for that experiment as the response. This dual approach helps evaluate how well the method detects perturbations consistently across both splicing factors and individual perturbation experiments.

As an additional metric, we computed the rank percentile of the perturbed splicing factor in each experiment. This score reflects how highly the true regulator ranks among all splicing factors given estimated activity, providing a simple and robust measure of performance.

Unless stated otherwise, all performance metrics were computed using held-out benchmark datasets.

### Robustness of empirical splicing factor networks

To evaluate the robustness of empirical splicing factor networks based on the likelihood scores of splicing factor→exon interactions, we pruned the targets of each splicing factor keeping the 40, 50, 60, 70, 80, 90, and 100 interactions with the highest likelihood. We then evaluated pruned versions of splicing factor networks by computing within- and across-sample recall as described above.

### Experimental validations of splicing factor activity estimation with diverse perturbations

#### Indisulam-induced degradation of RBM39 splicing factor protein

We downloaded the proteomic and transcriptomic data produced by Nijhuis *et al.*^23^ upon perturbation of IMR-32 neuroblastoma cells with either Indisulam (5 μM, 6 hours) or DMS vehicle control. Proteomic experiments had 3 biological replicates per condition while transcriptomic experiments had 1 biological replicate per condition. On the proteomics side, we obtained processed proteomic data in the form of LFQ values for each protein and sample (see Data Availability). Then, considering only measurements for splicing factors, we performed a statistical differential analysis of protein expression comparing Indisulam-treated to control samples through a T-test for each splicing factor using the function *ggpubr::compare_means(method=”t.test”)* and computed the average log2 fold changes between conditions. On the transcriptomics side, we processed raw RNA sequencing FASTQ files with *vast-tools* as described above to quantify exon inclusion (PSI) and gene expression (TPM). To estimate splicing factor activities with VIPER, we computed the splicing signatures of the Indisulam-treated cells by subtracting exon inclusion measured in control samples and ran VIPER with empirical splicing factor networks as explained above.

Finally, because the Nijhuis *et al.*^23^ dataset had a single biological replicate for transcriptomic data condition (see Supplementary Fig. 3a), we obtained RNA sequencing data from Lu *et al.*^24^, who measured the transcriptomic changes upon treatment with Indisulam (1 μM, 96 hours) compared with DMSO vehicle control with three biological replicates per condition for three cell lines (A375, MEL501, SKMEL239). After processing raw files with *vast-tools* as described above, we confirmed the changes in mRNA levels of RBM39 upon Indisulam treatment (see Supplementary Fig. 3b).

#### Combinatorial knockdown of splicing factors

For the four studies considered that present measured transcriptomic changes upon combinatorial knockdown of splicing factors^25–28^, we processed their raw RNA sequencing data with *vast-tools* as described above to obtain exon inclusion profiles. For each study, cell line, and condition, we obtained the splicing signatures required for VIPER inference of splicing factor activity by subtracting the average exon inclusion in unperturbed control samples from the corresponding perturbed samples. We estimated splicing factor activities with empirical splicing factor networks as described above. And, when applicable, we median-summarized splicing factor activities across biological replicates. This resulted in a table of predicted activities per condition and splicing factor in the empirical splicing factor networks.

#### Recapitulation of SF3b complex protein-protein interactions

We measured the shortest paths from each protein to each splicing factor in the SF3b complex within the STRINGDB protein-protein interaction network using the *networkx.shortest_path_length* function. Then, we summarized shortest path lengths to the SF3b complex level considering the minimal shortest path lengths among the splicing factors in the protein complex. We placed all interactions with a minimal shortest path length equal to or greater than 4 in the same bin.

We estimated splicing factor activities for six studies that measured how the transcriptome changes when perturbing cell lines with spliceosome drugs^19,65–69^. We processed raw RNA sequencing data with *vast-tools* as described above to obtain exon inclusion profiles. For each study, cell line, drug concentration, and incubation time, we obtained splicing signatures by subtracting the average exon inclusion in unperturbed vehicle controls from the corresponding perturbed condition. We then computed splicing factor activities with VIPER using empirical splicing factor networks as described above. Finally, when applicable, we summarized the estimated splicing factor activities across biological replicates in each condition by computing the median.

### Identification of recurrent splicing programs in cancer

We processed raw RNA sequencing samples from TCGA with *vast-tools* as described above to obtain exon inclusion and gene expression tables for all 33 types of cancer available. To identify recurrent splicing programs associated with cancer, for each type of cancer, we compared primary tumor samples to their healthy solid tissue normal counterparts provided each condition had at least 10 samples. Hence, we could only perform this comparison for 14 types of cancer.

#### Cancer splicing programs based on splicing factor activity

To perform differential activity analysis, we require predicting splicing factor activity in both tumor and healthy samples. We reasoned we could obtain splicing signatures for both conditions using the median of the other as a reference. Specifically, for each cancer cohort with sufficient data, we obtained splicing signatures for each tumoral sample by subtracting the median exon inclusion across healthy samples and we obtained splicing signatures for each healthy sample by subtracting the median exon inclusion across tumor samples. We used these signatures to compute VIPER splicing factor activities with empirical splicing factor networks as explained above. This resulted in a table of splicing factor activities for each tumor- and healthy-derived sample in each type of cancer. We performed differential activity analysis for each splicing factor by computing the median activity difference between tumor and healthy conditions and by statistically testing the significance of these differences through two-sided Wilcoxon rank sum tests using the function *scipy.stats.mannwhitneyu*. We considered as significant all those differences in splicing factor activity with an FDR (adjusted *p*-value) lower than 0.05. We quantified the recurrence of splicing factor activation and inactivation by counting in how many of the 14 types of cancers considered each splicing factor was differentially activated (oncogenic-like) or inactivated (tumor suppressor-like) in tumor samples. We assigned splicing factors to the oncogenic cancer splicing program if they were more frequently activated than inactivated in tumors with a threshold difference of 5. And, correspondingly, we assigned splicing factors to the tumor suppressor cancer splicing program if they were more frequently inactivated than activated in tumors with a threshold difference of 5.

#### Cancer splicing programs based on splicing factor gene expression

To identify cancer splicing programs based on gene expression, we used DESeq2^70^ to perform differential expression analysis. Starting with raw gene count data, we compared primary tumor samples to normal solid tissue samples for each of the 14 cancer cohorts included in our study.

For each splicing factor, we assessed whether its expression was significantly different (FDR < 0.05). We then counted the number of cancer cohorts where each splicing factor was consistently upregulated or downregulated. To define “oncogenic-like” and “tumor suppressor-like” programs, we required a splicing factor to be consistently upregulated in at least 5 cancer types to be classified as “oncogenic-like”. Similarly, splicing factors downregulated in at least 5 cancer cohorts were classified as tumor suppressor-like.

### Association sign of molecular measurements with patient outcome

For the 20 cancer cohorts in the TCGA with information regarding patient age and sex, and tumor staging, we studied the prognostic power of the splicing factors in recurrent cancer splicing programs identified either through splicing factor activities predicted with VIPER or through splicing factor mRNA levels by fitting Cox Proportional Hazards (Cox PH) models using the *survival::coxph* function that associate splicing factor activities or gene expression in tumor samples to patient clinical data (overall survival and survival status). We also considered age, sex and tumor staging metadata columns as covariates of the model. Fitted model coefficients inform how the increase of the variable of interest is associated with an increase (positive coefficient) or decrease (negative coefficient) in the risk of death.

For this analysis, we had to re-estimate splicing factor activities with VIPER as we aimed to study how splicing factor activities varied across tumors regardless of the molecular state of healthy counterparts. We obtained splicing signatures for each type of cancer by subtracting median exon inclusion across the tumor samples from each cancer cohort. Then, we computed splicing factor activities with VIPER using empirical splicing factor networks as described above.

### Overrepresentation analysis (ORA)

To assess gene set enrichment, we used the *clusterProfier::enricher* function, which performs statistical tests to identify significant overlaps between our gene lists and predefined pathways. We used ReactomeDB as our reference database.

For background (or “universe”) genes, we kept the default setting, which uses all genes annotated in ReactomeDB.

### Identification of exons in splicing programs associated with immune evasion

#### Analysis of melanoma patients treated with immune checkpoint blockade drugs

As mentioned in the results section, genes carrying exons in the tumor suppressor-like cancer splicing program were enriched in immune evasion-related terms. To study whether exons in genes belonging to these gene sets could play a role in response to ICB treatment, we processed raw RNA sequencing data with *vast-tools* as described above for the 109 tumor samples of cancer patients treated with ICB from the *Riaz et al.*^43^ study. We then classified each patient into a responder or non-responder considering their RECIST^42^ classification; CR (n=3) and PR (n=7) patients were considered as responders (n=10), and PD patients as non-responders (n=23). For each of the 608 exons in the 181 genes found in enriched immune invasion terms, we performed a differential splicing analysis comparing responders to non-responders. We computed exon inclusion differences between the groups by subtracting median exon inclusion in non-responders from median exon inclusion in responders and we tested these differences statistically through two-sided Wilcoxon rank sum tests using the function *ggpubr::compare_means(method=”wilcox.test”)*. We only reported as significant those exon inclusion differences with a nominal *p*-value < 0.05.

In addition, for the case of exon HsaEX1036341_SEC22B, we performed a Kaplan-Meier curve analysis to relate overall patient survival probability over time while stratifying by the inclusion of this exon. To find the best possible split in exon HsaEX1036341_SEC22B inclusion, we used the function *survminer::surv_cutpoint* with default parameters.

#### In vivo genome-wide CRISPR knockout screening data

We used data from the Dubrot *et al.*^71^ study to obtain the score quantifying the contribution of each gene to immune evasion in immunocompetent mice model treated with ICB compared with immuno-deficient mice. After downloading that data, we translated mouse gene names to human gene names using a table generated through BioMart (https://www.ensembl.org/info/data/biomart/index.html). Then, we multiplied the “Average Score” column by the “Sign” column to obtain signed scores that are informative of whether knocking out a given gene favors (positive) or hampers (negative) immune evasion and we kept only those signed scores measured for the condition “ICB vs NSG”. Finally, we only considered those genes whose exon was differentially spliced in the analysis above.

### Association of splicing factor activity with proliferative state marker MKI67

We processed raw RNA sequencing data from the CCLE with *vast-tools* as described above to quantify both exon inclusion and gene expression for the 1,015 cell lines with sufficient sequencing coverage. To estimate splicing factor activities with VIPER, we computed splicing signatures for each cell line by subtracting the median exon inclusion from the exon inclusion matrix. We predicted splicing factor activities with VIPER using empirical splicing factor networks as described above.

### Activities of cancer splicing programs during carcinogenesis

We processed raw RNA sequencing data from the Danielsson *et al.*^46^ study with *vast-tools* as described above. To estimate how splicing factor activities change at each stage of BJ fibroblast transformation, we computed the splicing signatures by subtracting average exon inclusion in unperturbed cells from transformed cells. We then predicted splicing factor activity with VIPER using empirical splicing factor networks as explained above.

### Proteomics and phosphoproteomics of primary fibroblasts during induced carcinogenesis

#### Cell culture

BJ fibroblasts from the Danielsson *et al.*^46^ study were cultured at 37 °C in a 5% (vol/vol) CO2 environment with Dulbecco’s modified Eagle’s medium (DMEM) supplemented with 10% fetal bovine serum (FBS). Each cell line –BJ primary, BJ immortalized (hTERT), BJ tumorigenic (hTERT + SV40), and BJ metastatic (hTERT + SV40 + RASG12V)– was cultured in two biological replicates to follow the experimental design in Danielsson *et al.*^46^.

#### Sample preparation

##### Cell harvesting

When reaching 90% of confluence, cells were washed with PBS and lysed in 100 µl of Urea 6 M, NH4HCO3 0.2 M. The samples were sonicated for 10 min at high position with on/off pulses of 30 seconds (Diagenode Bioruptor) and then centrifuged at 4°C for 10 min at 16000 g. Total protein in the extracts was determined by BCA assay (Pierce) and adjusted at 1 µg protein/µl.

For phosphoproteomic characterization, samples were treated with a phosphatase inhibitor by adding 100 µl of Urea 6 M, NH4HCO3 0.2 M, and PhosSTOP (1X) from Roche.

##### In-solution digestion

Samples (500 µg) were reduced with dithiothreitol (1500 nmol, 37 °C, 60 min) and alkylated in the dark with iodoacetamide (3000 nmol, 25 °C, 30 min). The resulting protein extract was first diluted to 2M urea with 200 mM ammonium bicarbonate for digestion with endoproteinase LysC (1:10 w:w, 37°C, over 6h, Wako, cat # 129-02541), and then diluted 2-fold with 200 mM ammonium bicarbonate for trypsin digestion (1:10 w:w, 37°C, over 8h, Promega cat # V5113).

After digestion, peptide mix was acidified with formic acid and desalted with a MicroSpin C18 column (The Nest Group, Inc).

##### Phosphoenrichment

Samples (490 µg of peptide mix) were enriched in phosphopeptides with the High-Select™ TiO2 Phosphopeptide Enrichment Kit (Thermo Scientific, cat # A32993).

#### Chromatographic and mass spectrometric analysis

Samples (proteome and phosphoproteome) were analyzed using an Orbitrap Astral mass spectrometer (Thermo Fisher Scientific) coupled to a Vanquish Neo liquid chromatographer (Thermo Fisher Scientific). Peptides were loaded directly onto the analytical column and were separated by reversed-phase chromatography using a 50 cm μPAC™ column (Thermo Scientific, cat # COL-NANO050NEOB), featuring a structured pillar array bed with a 180 µm bed width.

Chromatographic gradient was initiated with 96% buffer A and 4% buffer B at a flow rate of 750 nL/min. Over 1.9 minutes, the percentage of buffer B was increased to 10%. The flow rate was then reduced to 250 nL/min, and the percentage of buffer B was further increased to 22.5% over 23 minutes. Buffer A: 0.1% formic acid in water. Buffer B: 0.1% formic acid in 80% acetonitrile.

The mass spectrometer was operated in positive ionization mode with data-independent acquisition, with full MS scans over a mass range of m/z 380-980 with detection in the Orbitrap at a resolution of 240,000. In each cycle of data-independent acquisition, 300 windows of 2 Th were used to isolate and fragment all precursor ions from 380 to 980 m/z. A normalized collision energy of 25% was used for higher-energy collisional dissociation (HCD) fragmentation. The MS2 scan range was set from 150 to 2000 m/z with detection in the Astral with a maximum injection time of 3 ms.

Digested bovine serum albumin (New England Biolabs cat # P8108S) was analyzed between each sample to avoid sample carryover and to assure stability of the instrument, and QCloud^72^ has been used to control instrument longitudinal performance during the project.

#### Analysis of mass spectra

Acquired spectra were analyzed with DIA-NN (Neural networks and interference correction enable deep proteome coverage in high throughput) (v 2.1) using a predicted spectral library from the Swiss-Prot human database (as of June 2025, 20421 entries) plus a list^73^ of common contaminants.

For peptide identification, a precursor ion mass tolerance of 7 ppm was used for MS1 level, trypsin was chosen as the enzyme, and up to one missed cleavage was allowed.

For the proteome, oxidation of methionine and N-terminal protein acetylation were used as variable modifications, whereas carbamidomethylation on cysteines was set as a fixed modification.

For the phosphoproteome phosphorylation of serine, threonine, and tyrosine, oxidation of methionine and N-terminal protein acetylation were used as variable modifications, whereas carbamidomethylation on cysteines was set as a fixed modification.

The precursor and fragment ion m/z mass ranges were adjusted to 380-980 and 150-2000, respectively. False discovery rate (FDR) was set to a maximum of 1% at both the precursor and protein levels. For peptide quantification, match-between-runs was enabled, protein inference was set to ‘Protein names (from FASTA)’ with the ‘Peptidoforms’ option. The single-pass Neural Network machine learning model was used with the Quant UMS high precision quantification strategy. Default settings were used for the other parameters.

The pg-matrix or pr-matrix output (proteome and phosphoproteome, respectively) from DIA-NN was subsequently used to analyze differential expression using FragPipe Analyst^74^ (v1.14), employing all default settings except for imputation, which was set to "No imputation”.

We summarized the proteomic data at the protein level, rather than the peptide level. For proteins with multiple detected peptides, we assigned the maximum LFQ (Label-Free Quantification) value among their peptides (referred to as “max LFQ”). To normalize the count matrices for both proteins and phosphopeptides, we adjusted for sample-specific depth by dividing each count by the total number of counts observed in its respective sample. We then scaled values to counts per million by multiplying by 10^6^. After normalization, we applied log2(count + 1) transformation to stabilize variance. In total, we detected 10,349 unique proteins and 60,707 phosphopeptides, corresponding to 6,963 distinct proteins.

### Identifying putative multi-omic drivers of cancer splicing program regulation using regularized linear models

In the study by Danielsson *et al.*^46^, we observed a coordinated shift in activity of cancer splicing programs during carcinogenesis: oncogenic-like splicing factors become more active, while tumor suppressor-like splicing factors lose activity at the tumorigenic stage.

To investigate the molecular drivers of this shift, we integrated multiple omic layers for splicing factors:

- mRNA expression and splicing event usage (exons, introns, alternative splice sites) from Danielsson *et al.*^46^
- Protein abundance and phosphorylation levels from the current study

For each omic feature, we calculated the difference relative to the average of control samples (primary BJ fibroblasts). We removed features with missing or constant values across samples, resulting in 7,575 multi-omic features: mRNA expression (n = 484), exon inclusion (n = 1,049), intron retention (n = 1,970), alternative donor sites (n = 438), alternative acceptor sites (n = 700), protein abundance (n = 444), phosphopeptide abundance (n = 2,490).

To summarize the relative activation of cancer splicing factor programs, we computed the difference between the median activity of oncogenic-like splicing factors and tumor suppressor-like splicing factors in each condition.

To link these molecular changes with shifts in splicing program activity across carcinogenic stages (immortalized, tumorigenic, and metastatic), we applied regularized linear regression using the *glmnet* package^75^. With only six observations (three stages in biological duplicates), we chose lasso regression (alpha = 1) and elastic net (alpha = 0.5) to ensure robustness and feature selection.

We interpreted the models by examining non-zero coefficients, which identify omic features that explain the observed changes in splicing program activity during cancer progression.

## DATA AVAILABILITY

### In this study

The raw proteomics and phosphoproteomics data have been deposited in the PRIDE^76^ repository via ProteomeXchange with identifier PXD066623.

### Intermediate files generated from data analyses

Intermediate files generated throughout this study are available in this Figshare repository: https://doi.org/10.6084/m9.figshare.27835518

### Previously published datasets used

*List of splicing factors from Papasaikas et al.^18^*
Supplementary Table 2 in the publication.

*List of splicing factors from Hegele et al.^21^*
Supplementary Table 1 in the publication.

*List of splicing factors from Seiler et al.^19^*
Supplementary Table 1 in the publication.

*List of splicing factors from Rogalska et al.^8^*
Supplementary Table 1 in the publication.

*List of splicing factors from Head et al.^20^*
Supplementary Table 1 in the publication.

*RNA sequencing data of splicing factor perturbations from ENCORE project*
We downloaded raw FASTQ files from the ENCORE website for all shRNA and CRISPR experiments targeting RNA binding proteins: https://www.encodeproject.org/encore-matrix/?type=Experiment&status=released&internal_tags=ENCORE.

*RNA sequencing data of splicing factor perturbations from ENA*
We downloaded raw FASTQ files from the ENA website listed in Supplementary Table 2 in this manuscript.

*POSTAR3 CLIP data to construct CLIP-based splicing factor networks* http://postar.ncrnalab.org/

*Observational RNA sequencing data from human samples from diverse molecular contexts*
We downloaded raw FASTQ files for Cardoso-Moreira *et al.^60^* through the ENA website.

*Perturbing RBM39 with Indisulam*
For the Nijhuis *et al.^23^* study, we downloaded raw FASTQ files from the ENA website (PRJNA673205) and processed LFQ proteomics data from the PRIDE website (PXD022164). Metadata for proteomics experiments were kindly provided by the authors.
For the Lu *et al.^24^* study, we downloaded raw FASTQ files from the ENA website (PRJNA683080).

*Combinatorial knockdowns of splicing factors*
We downloaded raw FASTQ files for four different studies from the ENA website: PRJNA223244^28^, PRJNA498529^27^, PRJNA587741^26^, and PRJNA321560^25^.

*Protein-protein interaction network*
We downloaded STRING DB’s human protein interaction network from their webpage^77^. https://stringdb-static.org/download/protein.links.full.v11.5/9606.protein.links.full.v11.5.txt.gz

*List of genes in SF3b complex* https://signor.uniroma2.it/relation_result.php?id=SIGNOR-C442

*List of genes in U2snRNP complex* https://signor.uniroma2.it/relation_result.php?id=SIGNOR-C479

*Perturbing SF3b complex reaction step with splicing drugs*
We downloaded raw FASTQ files for six different studies from the ENA website: PRJNA371421^30^, PRJNA685790^65^, PRJNA662572^66^, PRJNA380104^67^, PRJNA354957^68^, and PRJNA292827^69^

*Splicing event information*
We downloaded information and their predicted protein impact of splicing events from VastDB. https://vastdb.crg.eu/wiki/Downloads#Homo_sapiens_.28hg38.29

*Molecular and clinical data from The Cancer Genome Atlas (TCGA)*
We used the GDC portal to download the clinical, survival, and raw RNA-seq data.

*SpliceosomeDB complexes*
We obtained annotations of splicing factors to splicing reaction complexes from the SpliceosomeDB website. http://spliceosomedb.ucsc.edu/

*RBP domain families*
We obtained annotations of splicing factors to RBP domain families from the RBPDB website. http://rbpdb.ccbr.utoronto.ca/

*Reactome pathways processed by MSigDB*
We obtained ReactomeDB pathways from the MSigDB website. https://www.gsea-msigdb.org/gsea/msigdb/human/genesets.jsp?collection=CP:REACTOME

*Immune evasion CRISPR screen from Dubrot et al.^71^*
We obtained the immune evasion score for each gene from Supplementary Table 13 in the publication. https://static-content.springer.com/esm/art%3A10.1038%2Fs41590-022-01315-x/MediaObjects/41590_2022_1315_MOESM2_ESM.xlsx

*Human to mouse gene name translations*
We obtained tables of human-to-mouse gene name translations from BioMart.

*Data from patients treated with ICB from Riaz et al.^43^*
We obtained raw FASTQ files from the ENA website (PRJNA356761).
We obtained patient clinical metadata from Supplementary Table 2 in the publication.

*Molecular data from the Cancer Cell Line Encyclopedia (CCLE)*
We downloaded raw RNA-seq .fastq files for the cancer cell lines in the CCLE from the ENA website (PRJNA523380).
We downloaded cell line metadata (“sample_info.csv”) and processed mutation mapping across CCLE cell lines (“CCLE_mutations.csv”) from DepMap’s figshare repository (download link: https://ndownloader.figshare.com/articles/13681534/versions/1).

*RNA sequencing data of carcinogenesis from Danielsson et al.^46^*
We downloaded raw FASTQ files from the ENA website (PRJNA193487).

*DEMETER2 gene dependencies from DepMap* https://ndownloader.figshare.com/articles/6025238/versions/6

*CCLE cell line metadata*
https://ndownloader.figshare.com/articles/13681534/versions/1

## CODE AVAILABILITY

To improve the reproducibility of our results, the full data analysis pipeline of this manuscript will be made publicly available at https://github.com/MiqG/publication_viper_splicing and the corresponding scripts to run VIPER with the splicing factor→exon networks identified in this study at https://github.com/MiqG/viper_splicing.

---

**Extended Data Figure 1.**
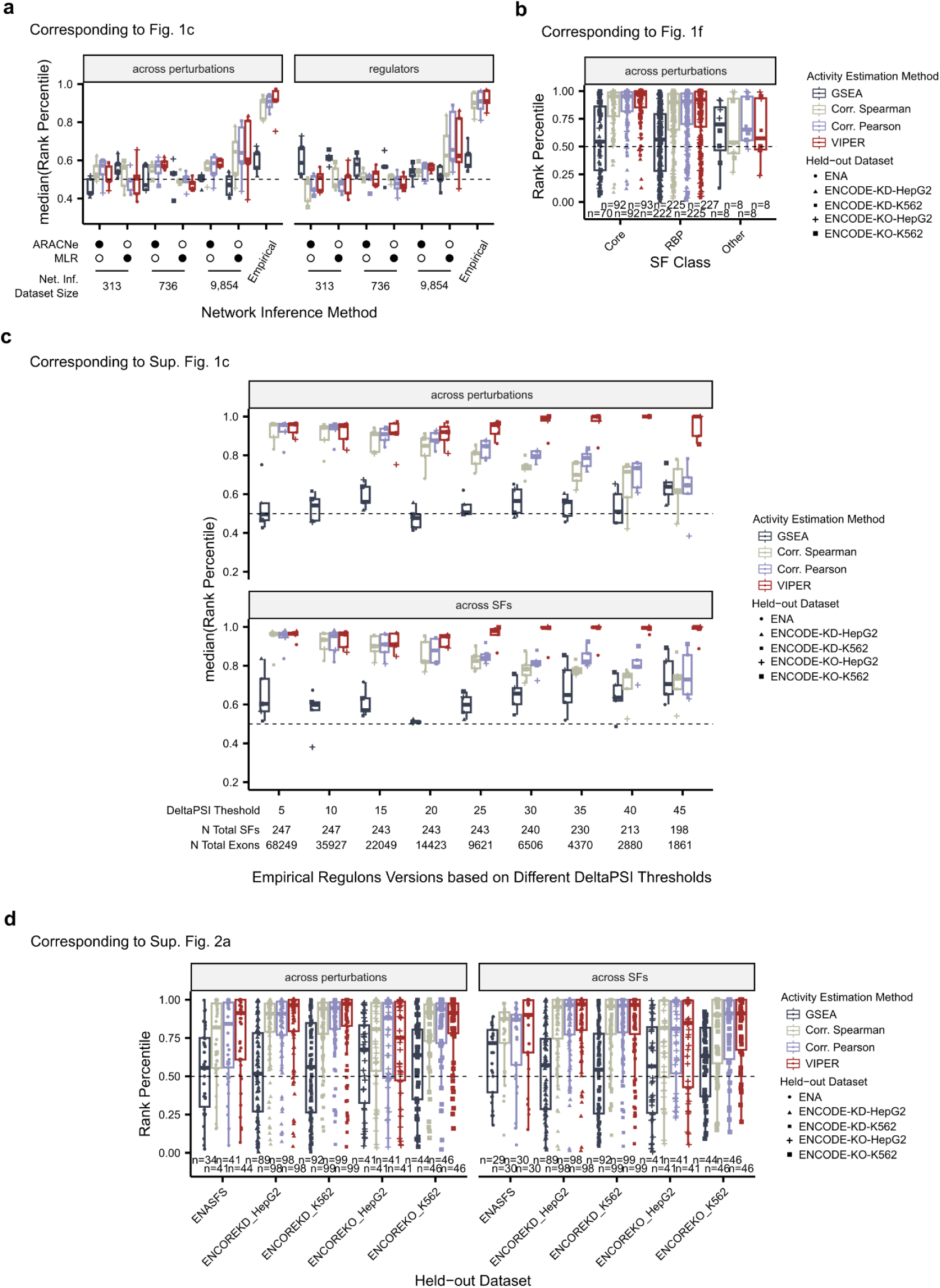
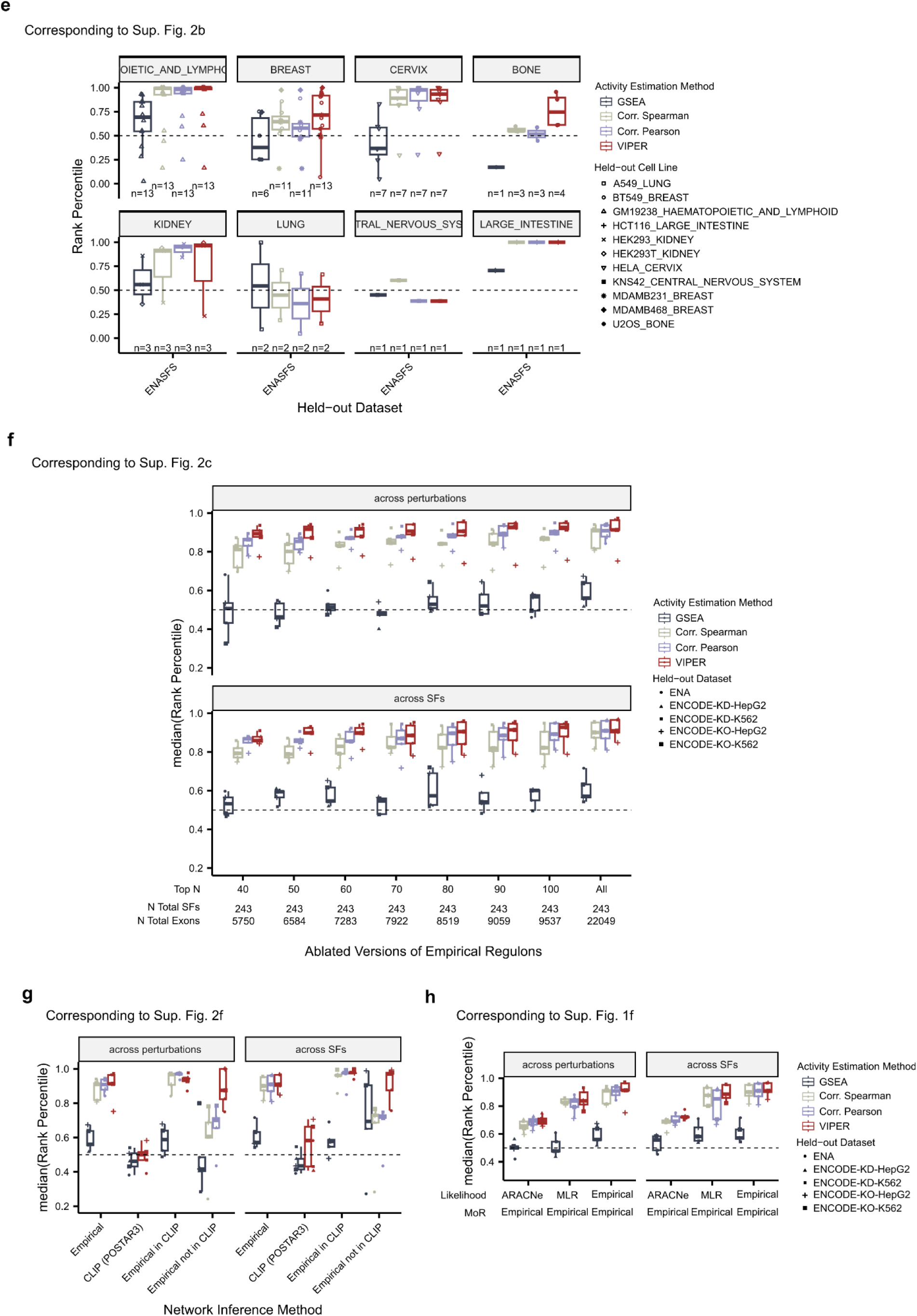
Reproducing splicing factor activity benchmarks with rank percentiles as an alternative metric to ROC AUCs does not alter conclusions. Each panel includes a box plot with rank percentile scores (Y-axis) that correspond to of ROC-AUC-based evaluations when comparing (**a**) network inference methods; (**b**) classes of splicing factors; (**c**) exon thresholds to define empirical splicing factors networks; (**d**) performance on individual experiments for each held-out benchmark dataset; (**e**) evaluation considering the cell line lineages of ENA held-out dataset; (**f**) network ablation of weakest splicing factor→exon interactions; (**g**) direct (CLIP interaction) vs indirect (no CLIP interaction) empirical networks; (**h**) combinations of empirical mode of regulation with computationally-inferred splicing factor→exon interactions. At the top of each panel we indicate the corresponding ROC-AUC-based panel.

## ACKNOWLEDGEMENTS

We thank William C. Hahn, Annica Gad, Emma Lundberg, and Staffan Strömblad for kindly sharing the BJ fibroblasts and its derivatives with us. And, Attila Biçer and Matilde D’Angelo for their help in culturing BJ fibroblast cell lines. We would like to thank Luca Zanella, Lucas ZhongMing Hu, Alessandro Vasciaveo, Alec Wang for their support in the reimplementation of VIPER. We would like to thank Sophie Bonnal, Malgorzata Ewa Rogalska, Juan Valcárcel, Manuel Irimia, Yamile Ana, Javier Gonzalez-de-Miguel and Xavier Hernandez-Alias for the insightful discussions regarding the elucidation of splicing factor programs. Also, we would like to thank Hector C Keun, Clare Eckold and Anke Nijhuis for kindly sharing proteomics sample metadata and raw RNA sequencing data from cells treated with indisulam. We acknowledge support of the Spanish Ministry of Science and Innovation through the Centro de Excelencia Severo Ochoa (CEX2020-001049-S, MCIN/AEI /10.13039/501100011033), and the Generalitat de Catalunya through the CERCA programme, and to the EMBL partnership. We are grateful to the CRG Core Technologies Programme for their support and assistance in this work. The proteomics analyses were performed in the CRG/UPF Proteomics Unit, which is part of the Spanish National Infrastructure for Omics Technologies (ICTS OmicsTech). The results shown here are in part based upon data generated by the TCGA Research Network (https://www.cancer.gov/tcga).

## AUTHOR INFORMATION

Conceptualization: MA-G, SM-V, AC, LS; Methodology: MA-G, CS-M, DFM, SM-V, AC, LS; Software: MA-G, DFM, CZ; Validation: MA-G, CS-M, DFM, SM-V, AC, LS; Formal analysis: MA-G, DFM, SM-V, AC, LS; Investigation: MA-G, CS-M; Resources: AC, LS; Data Curation: MA-G; Writing - original draft: MA-G; Writing - review & editing: MA-G, DFM, SM-V, AC, LS; Visualization: MA-G; Supervision: SM-V, AC, LS; Project administration: MA-G; Funding acquisition: MA-G, CZ, AC, LS. All authors read and approved the final manuscript.

