## Supplementary Information for "Exon inclusion signatures enable accurate estimation of splicing factor activity"

#### SUPPLEMENTARY NOTES

##### Supplementary Note 1. Benchmark datasets for splicing factor network inference and activity estimation

To evaluate how different splicing factor network inference approaches work in combination with different activity estimation methods, we developed a set of benchmark datasets. We generated a consensus list of 509 splicing factors documented in landmark literature (Supplementary Fig. 1a; Supplementary Table 1 and 2)[1–5]. We then compiled and uniformly processed bulk RNA-seq data from the ENCODE and ENA databases, resulting in a total of 474 experiments that perturb 248 distinct splicing factors across 39 cell lines and 54 independent studies (Supplementary Fig. 1b). For each experiment, we computed exon inclusion signatures (delta PSI) by comparing perturbed versus unperturbed exon inclusion profiles and recorded the splicing factor that was perturbed.

The splicing signatures were divided into five benchmark datasets based on their source databases: ENA (n=161), ENCODE-KD-HepG2 (n=104), ENCODE-KO-HepG2 (n=44), ENCODE-KD-K562 (n=104), and ENCODE-KO-K562 (n=61). These benchmark datasets enable computing performance metrics such as the area under the receiver operating curve (ROC-AUC) to quantify how well the predicted splicing factor activities recapitulate the splicing factor responsible for the observed splicing signatures. ROC-AUCs can be calculated along two axes for each dataset: across splicing factors and across perturbations. These axes indicate the predictive power of the estimated splicing factor activities in identifying which experiments perturbed a specific splicing factor and which splicing factor was perturbed in each experiment, respectively (Fig. 2b).

##### Supplementary Note 2. Empirical splicing factor networks combined with VIPER outperform computational ones in activity estimation

Traditionally, splicing factor networks have been reverse-engineered computationally by assessing pairwise associations between splicing factor expression and exon inclusion across multiple samples[6]. To replicate this, we adapted ARACNe[7,8] –an algorithm that calculates pairwise mutual information between splicing factor expression (TPM) and exon inclusion (PSI)– to infer splicing factor networks using RNA-seq datasets with increasing sample sizes ($n_{CardosoMoreira2020}$ = 313, $n_{PANCAN\text{-}STN}$ = 736, and $n_{PANCAN\text{-}PT}$ = 9,854) (see Methods). This resulted in three computationally inferred splicing factor networks for the 509 annotated splicing factors. These networks consist of splicing factor→exon interactions with their associated likelihood (from 0 to 1) and mode of regulation (positive or negative).

We evaluated network quality for splicing factor activity estimation using VIPER and three additional baseline methods: normalized enrichment scores (NES) from gene set enrichment analysis (GSEA), and Spearman and Pearson correlations. Each method interprets splicing factor networks differently. GSEA, a widespread approach, uses interactions in an ontology (here, splicing factor networks) to compute NES on exon inclusion signatures, but it does not consider likelihoods or modes of regulation. In correlation-based methods, we incorporated likelihood and mode of regulation by multiplying them into an interaction summary value and computing a correlation coefficient (here, activity) between exon inclusion signatures and each splicing factor's target exons interaction summary values. Finally, VIPER combines aspects of both approaches by using interaction likelihoods and modes of regulation in an enriched rank-based analysis (aREA) to estimate splicing factor activity from exon inclusion signatures.

The ARACNe-based networks resulted in splicing factor activity estimates that were close to random, with slightly better performance when more samples were used, especially when combined with VIPER (Fig. 2c). This indicates that pairwise statistical dependencies between splicing factor expression and exon inclusion are not sufficient for effective network inference.

To address this limitation, we inferred splicing factor networks using multivariate linear regression (MLR) models, which simultaneously associate the expression of all splicing factors with exon inclusion, potentially accounting for combinatorial regulation (see Methods). However, this computational approach also performed poorly, yielding near-random ROC-AUC distributions in most cases, except for the largest dataset, where performance improved slightly (ROC-AUC ~ 0.6) (Fig. 2c).

Given the limitations of computational methods, we turned to empirical splicing factor networks. These networks designate exons as targets if their inclusion levels change by more than a defined threshold following splicing factor perturbation. Although such target exons may result from both direct and indirect interactions, they are functionally linked to their respective splicing factor's activity. We based empirical networks on perturbation experiments used to create benchmark datasets, cataloging exon inclusion changes associated with 248 of the 509 annotated splicing factors. Here, we set the magnitude of delta PSI as the interaction likelihood and the sign as the mode of regulation. To select an optimal delta PSI threshold, we generated and evaluated nine networks across a range of delta PSI cutoffs (from 5 to 45% > |delta PSI|) (Supplementary Fig. 1c). To avoid overfitting, we excluded interactions derived from the benchmark dataset used for evaluation.

We found using a 15% delta PSI threshold produced networks that, when combined with VIPER, achieved highly accurate activity estimations (median ROC-AUC = 0.91 across both axes) while limiting the number of target exons (n = 22,049) (Supplementary Fig. 1c). These empirical networks outperformed both computational networks and other activity estimation methods, indicating that combining empirical splicing factor networks with VIPER provides the best approach for accurate splicing factor activity estimation (Fig. 2c).

##### Supplementary Note 3. Poor mode of regulation prediction limits the performance of computational networks

Given the quality of predictions based on the perturbation-derived empirical splicing factor networks, we wondered what may be causing the poor performance of computational approaches. For this benchmark, we used computational splicing factor networks inferred from the largest dataset ($n_{PANCAN-PT}$ = 9,854) as they showed the best performance. Splicing factor network interactions are assigned both a likelihood score (from 0 to 1) and a mode of regulation (positive or negative) (Fig. 2a). Computational methods successfully assigned higher likelihood to splicing factor→exon interactions that were observed in perturbation experiments, suggesting this may not be the source of poor performance (Supplementary Fig. 1d). Instead, the mode of regulation, which predicts whether an exon is included or excluded, was inferred almost randomly, with a prediction error close to 50% (Supplementary Fig. 1e).

To validate this, we tested whether combining ARACNe- or MLR-based likelihoods with experimentally derived modes of regulation would improve performance. Indeed, this approach boosted ROC-AUC values, bringing them closer to those achieved by empirical networks (Supplementary Fig. 1f).

These findings suggest that computational methods struggle to accurately assess exon inclusion versus exon exclusion when splicing factor activity changes, leaving room for future improvement.

**Supplementary Note 4. Stress testing splicing factor activity estimation with empirical networks**

Despite promising results, estimating splicing factor activity using empirical networks with VIPER may still be affected by technical and biological artifacts.

From a technical standpoint, splicing changes are often highly context-specific[9], which could lead to batch effects if empirical splicing factor networks are derived from different studies or cell lines. To assess robustness across studies, we evaluated held-hout benchmark datasets independently. VIPER combined with empirical networks consistently achieved the highest ROC-AUC distributions, indicating resilience to batch effects (Supplementary Fig. 2a). As an extreme example, VIPER with ENCODE-derived networks (from 2 cell lines) achieved median ROC-AUC values of 0.90 (across splicing factors) and 0.91 (across perturbations) on the ENA benchmark dataset (37 different cell lines). Only the ENCODE-KO-HepG2 dataset using Pearson outperformed this result. To further test robustness across cellular contexts, we constructed empirical networks from ENCODE perturbations and evaluated performance on ENA cell lines from various tissue origins. In all tissue lineages represented by at least two cell lines, VIPER achieved higher ROC-AUC values than any other method applied to the same network (Supplementary Fig. 2b). Altogether, this suggests that combining perturbation-based networks with VIPER provides robust splicing factor activity estimation across independent datasets and cellular contexts.

As an additional robustness check, we repeated all benchmarking analyses using an alternative performance metric: the rank percentile of the perturbed splicing factor among all factors, rather than ROC-AUC. This approach assesses whether the perturbed factor ranks near the top based on activity estimates. The results remained consistent with our original conclusions (Extended Data Fig. 1).

We further validated robustness by retaining only the top interactions in empirical networks based on likelihood. Reducing target exons by up to 73.9% did not substantially alter evaluation metrics for VIPER-assessed activities, mirroring findings from transcription factor activity benchmarks using this method (Supplementary Fig. 2c).

Biologically, cross-regulation among splicing factors may impact activity estimation. Overall, considering the ~140,000 annotated exons in VastDB, empirical splicing factor networks show low connectivity as each exon is regulated by a median of two splicing factors, and each splicing factor regulates a median of 300 exons (Fig. 2d). This indicates splicing changes from each splicing factor perturbation are highly specific, enhancing the accuracy of the resulting splicing factor networks.

Given the functional differences between splicing factor classes, we examined how connectivity in empirical networks correlates with their predictive power. Core factors are more connected than RNA-binding proteins (RBPs) or other splicing factors (Fig. 2e). Similarly, activity estimates for core factors reach higher ROC-AUC scores than the rest (Fig. 2f). This suggests that the exon repertoire of core splicing factors is more stable across conditions than those of RBPs or other splicing factors, which are expected to exhibit more context-dependent regulation[10].

**Supplementary Note 5. Indirect interactions dominate empirical networks, with VIPER ensuring robust activity estimation**

A possible limitation of our perturbation-based empirical splicing factor networks is that direct splicing factor→exon interactions are indistinguishable from indirect ones, potentially affecting performance. For some splicing factors, we can distinguish direct from indirect splicing factor→exon interactions. Crosslinking and immunoprecipitation (CLIP) assays identify RNA sequences physically bound by RBPs, providing direct interaction data. Since most splicing factors are RBPs, we explored whether identifying direct interactions in empirical splicing factor networks using CLIP data influences performance. Using POSTAR3[11], we generated CLIP-based networks for the 97 RBP splicing factors found. We included all splicing factor→exon interactions where CLIP peaks fell within 500 base pairs (bp) of an exon (Supplementary Fig. 2d). Only 11,626 of the 1,891,625 CLIP interactions overlapped with the 121,763 perturbation-based interactions, suggesting that most direct RBP-RNA interactions do not cause strong exon inclusion changes and that most changes in exon inclusion caused by splicing factor perturbations are likely due to indirect interactions (Supplementary Fig. 2e).

In terms of performance, CLIP-based networks alone produced near-random ROC-AUC scores. But, empirical networks containing only CLIP-verified (direct) interactions showed comparable performance to the original empirical networks using either correlation-based or VIPER for activity estimation (Supplementary Fig. 2f). Empirical networks with only non-CLIP-verified (indirect) interactions showed more variable ROC-AUC scores. Of note, only VIPER maintained median ROC-AUC scores of 0.8 or above when estimating activity from empirical networks of indirect interactions (Supplementary Fig. 2f). These findings confirm that, when possible, identifying direct targets can enhance activity estimation, and demonstrate VIPER's robustness in accurately estimating splicing factor activity, even with empirical networks containing only indirect interactions.

#### SUPPLEMENTARY FIGURE LEGENDS

##### Supplementary Figure 1. Evaluation of methods to reverse engineer splicing factor networks.

(**a**) UpSet plot illustrating the overlaps between the lists of splicing factors collected from the literature to obtain our consensus list of splicing factors.

(**b**) Overview of the data processed from publicly available repositories to evaluate the ability of inferred splicing factor networks to estimate splicing factor activities with VIPER and used to generate the empirical splicing factor network.

(**c**) Evaluation metrics of empirical splicing factor networks defined with different thresholds for absolute delta PSI.

(**d**) Likelihood scores from ARACNe and MLR match empirical interactions. Distributions of splicing factor→exon likelihood scores in ARACNe- or MLR-derived networks for experimentally observed vs. unobserved interactions. Top, two-sided Wilcoxon rank sum p-value. Bottom, interaction counts per category.

(**e**) MoR inferred by ARACNe and MLR approximate random (0.5). Proportion of correctly inferred positive and negative modes of regulation (MoR) interactions by ARANCe and MLR algorithms compared to experimental data.

(**f**) Combining ARACNe or MLR likelihoods with empirical MoR restores predictive power. Across perturbations and across splicing factors, ROC-AUCs for network versions using computational likelihoods with empirical MoR signs.

In box and whisker plots in panels (**c**), (**d**), and (**f**), the median is marked by a horizontal line, with the first and third quartiles as box edges. Whiskers extend up to 1.5 times the interquartile range, and individual outliers are plotted beyond.

##### Supplementary Figure 2. Continued evaluation of methods to reverse engineer splicing factor networks.

(**a**) Evaluation metrics of empirical splicing factor networks defined with four of the five benchmark datasets and evaluated on the fifth one.

(**b**) Evaluation metrics of empirical splicing factor networks derived from ENCODE perturbation datasets and evaluated on ENA cell lines of different lineages.

(**c**) Robustness analysis of the selected empirical splicing factor network (threshold delta PSI = |delta PSI|) consisting of keeping only top K interactions for each splicing factor sorted by splicing factor→exon interaction likelihood, or delta PSI magnitude.

(**d**) Distributions of distances between mapped CLIP peaks and target exons to generate CLIP-based splicing factor networks.

(**e**) Venn diagrams of the overlaps between the splicing factors of the empirical splicing factor networks and their target interactions.

(**f**) Evaluation of CLIP-based splicing factor networks compared to empirical networks. “Empirical in CLIP” are networks made of the overlapping interactions between perturbation-based networks and CLIP-based networks; direct empirical interactions. “Empirical not in CLIP” are empirical networks without CLIP-based interaction evidence; indirect empirical interactions. Note that correlation-based activities cannot be computed for CLIP-based networks as all likelihood and mode of regulation values are constant.

In box and whisker plots in panels (**a**), (**b**), (**c**), and (**f**), the median is marked by a horizontal line, with the first and third quartiles as box edges. Whiskers extend up to 1.5 times the interquartile range, and individual outliers are plotted beyond.

##### Supplementary Figure 3. Changes in transcriptomic gene expression upon treatment with Indisulam.

(**a**) mRNA levels of RBM39 treated with either Indisulam (5 μM, 6 hours) or DMSO vehicle control in the IMR-32 cell line from study Nijhuis *et al.*[12]. Top, corresponding gene expression values. Note this corresponds to a single biological replicate.

(**b**) mRNA levels of RBM39 of A375, MEL501, and SKMEL239 cell lines treated with either Indisulam (1 μM, 96 hours) or DMSO vehicle control from study Lu *et al.*[13]. Top, p-value based on two-sided Wilcoxon rank sum tests comparing the mRNA levels of cell lines in each condition.

In box and whiskers plots in panel (**b**), the median is marked by a horizontal line, with the first and third quartiles as box edges. Whiskers extend up to 1.5 times the interquartile range, and individual outliers are plotted beyond.

##### Supplementary Figure 4. Splicing programs defined by gene expression lack cancer driver-like behavior.

(**a**) Differential activation status of splicing factors (FDR < 0.05, two-sided Wilcoxon rank sum test) across cancer cohorts.

(**b**) Distributions of cancer cohorts in which each splicing factor is associated with high or low death risk, based on thresholds for defining recurrent activation in cancer splicing programs. Top, p-value based on two-sided Wilcoxon rank sum tests. Bottom, splicing factor count in resulting each program.

(**c**) Relationship between differential splicing factor protein activities and splicing factor gene expression fold-changes computed across 14 different types of cancer from TCGA comparing tumor samples to solid tissue normal (healthy) samples. Top, the Pearson correlation coefficient ($R$) and p-value ($p$).

(**d**) Frequency of differentially (FDR < 0.05) upregulated (top) and downregulated (bottom) splicing factors. We highlighted the top 5 splicing factors that were recurrently activated and inactivated in Fig. 3c. Horizontal dashed lines correspond to the selected recurrency threshold to be considered a splicing factor in either cancer splicing program. On top of the dashed line, the number of splicing factors selected as part of each gene expression-based cancer splicing program.

(**e**) Distributions of the number of cancer cohorts that each splicing factor expression is associated with a high or low risk of death in gene expression-based cancer splicing programs defined with recurrently differentially expressed splicing factors. Color, cancer splicing program. Top, p-value based on two-sided Wilcoxon rank sum tests. Bottom, number of splicing factors in the corresponding gene expression-based cancer splicing program.

(**f**) Distributions of the number of cancer cohorts that each splicing factor is associated with a high or low risk of death considering different thresholds of differential expression recurrence to define cancer splicing programs. Color, cancer splicing program. Top, p-value based on two-sided Wilcoxon rank sum tests. Bottom, number of splicing factors in the corresponding cancer splicing program.

In box and whisker plots in panels (**b**), (**e**), and (**f**), the median is marked by a horizontal line, with the first and third quartiles as box edges. Whiskers extend up to 1.5 times the interquartile range, and individual outliers are plotted beyond.

#### Supplementary Figure 5. Functional insights and dependencies of cancer splicing programs in carcinogenesis and immune response.

(**a**) The 10 most significantly enriched ReactomeDB terms (FDR < 0.05, ORA), ranked by gene set size, for the target genes of splicing factors PRPF8, SRSF1, and SF3B3. These factors are putative regulators of exon HsaEX1036341 in SEC22B.

(**b**) Distributions of Pearson correlations between proliferative state marker MKI67 and median splicing factor activities of each cancer splicing program. We considered the 22 cancer types with more than 10 cell lines in the database. Top, *p*-value based on two-sided Wilcoxon rank sum test.

(**c**) DEMETER2[14] gene dependency scores for splicing factors in each cancer splicing program, stratified by cell type and essentiality status of the splicing factor. Top, p-value based on Wilcoxon rank sum tests.

In box and whisker plots in panels (**b**), (**c**), and (**d**), the median is marked by a horizontal line, with the first and third quartiles as box edges. Whiskers extend up to 1.5 times the interquartile range, and individual outliers are plotted beyond.

#### Supplementary Figure 6. Multi-omic profiling of splicing factors across stages of carcinogenesis.

(**a-b**) Top panels, distributions of log-fold changes in mRNA expression, protein abundance, and phosphopeptide levels for splicing factors, comparing control BJ fibroblasts to the three carcinogenesis stages (immortalized, tumorigenic, metastatic). Bottom panels, corresponding differences in splicing event inclusion (“DeltaPSI”) for exons, introns, and alternative splice sites. In (**b**), the same distributions are shown as absolute values to highlight the overall magnitude of molecular changes. P-values above the distributions refer to two-sided Wilcoxon rank sum tests comparing each group of splicing factors against the “Non-driver” reference set. Values below the distributions refer to the number of splicing factor features that constitute the distribution.

(**c**) non-zero coefficients from regularized linear models linking splicing factor multi-omic features (expression, splicing, protein, phosphorylation) to shifts in cancer splicing program activity during carcinogenesis in the BJ fibroblast model. These associations suggest which features are putative drivers of the coordinated regulation of cancer splicing factor programs.

In box and whisker plots in panels (a) and (b), the median is marked by a horizontal line, with the first and third quartiles as box edges. Whiskers extend up to 1.5 times the interquartile range, and individual outliers are plotted beyond.

**Supplementary Figure 7. Parameter sensitivity analysis for PSI imputation and network inference with ARACNe**

(**a**) Evaluation of KNNImputer for PSI imputation with different k values. Distributions of real and imputed PSI values for 5 randomly selected samples from the CardosoMoreira2020 exon inclusion PSI table. “Synthetically Missing” PSI values were artificially set to NA for imputation evaluation. “Real” PSI values, left unaltered, are shown for comparison.

(**b**) Impact of “max_targets” on ARACNe inference. ROC AUC distributions for computational splicing factor networks inferred with ARACNe using different “max_targets” parameters. Networks were inferred using the CardosoMoreira2020 dataset. In box and whisker plots, the median is marked by a horizontal line, with the first and third quartiles as box edges. Whiskers extend up to 1.5 times the interquartile range, and individual outliers are plotted beyond.

#### SUPPLEMENTARY FIGURES

##### Supplementary Figure 1a-c

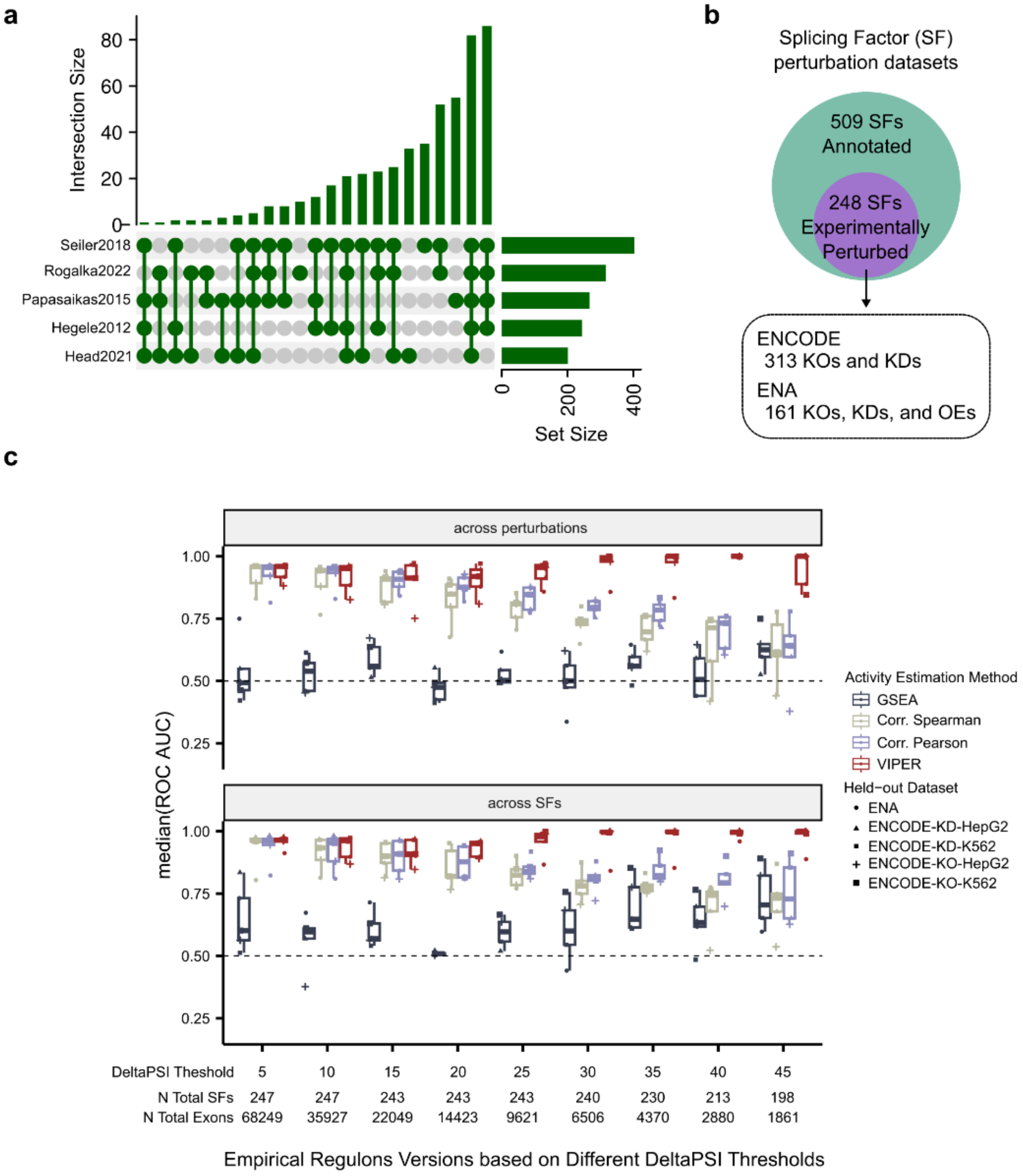

**Supplementary Figure 1d-f**

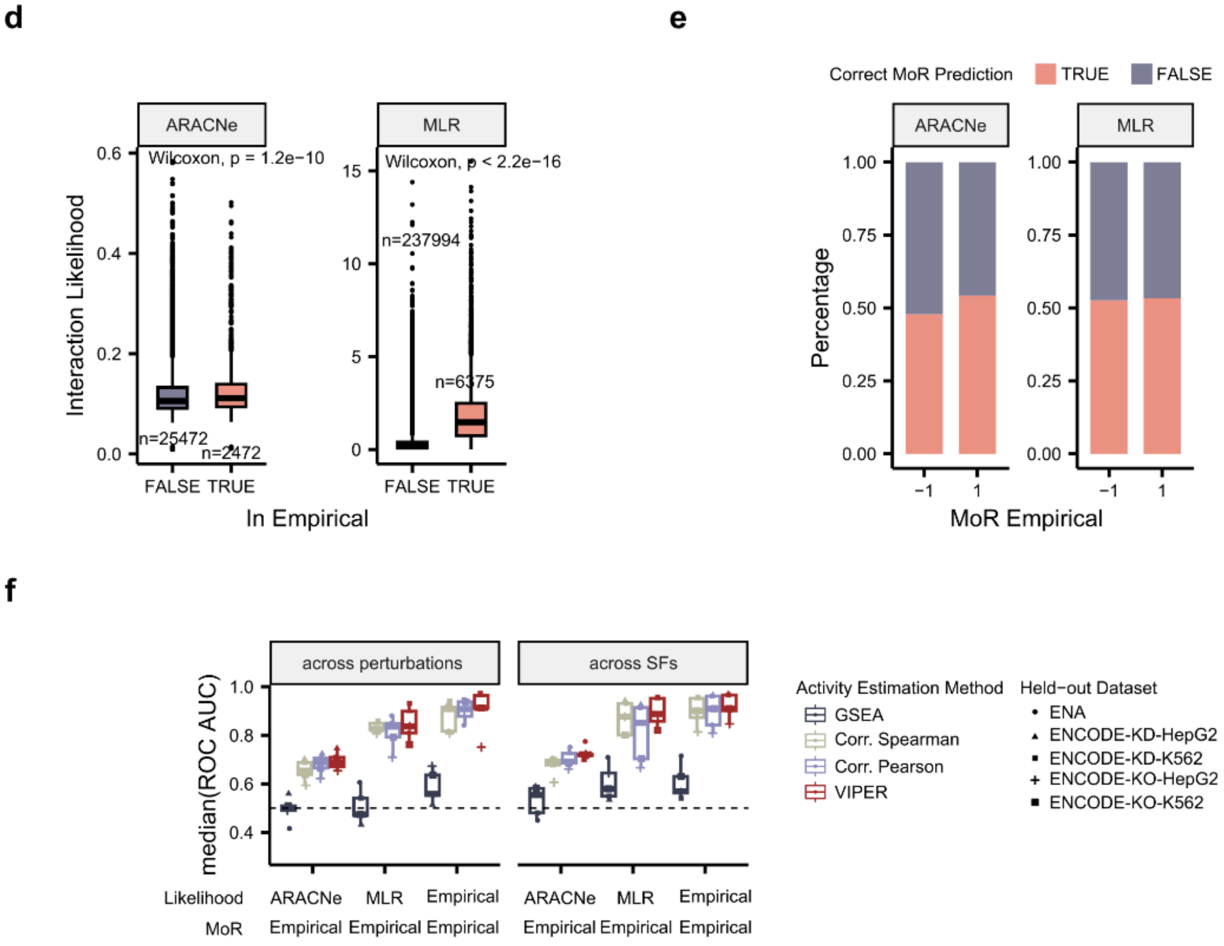

### Supplementary Figure 2a-c

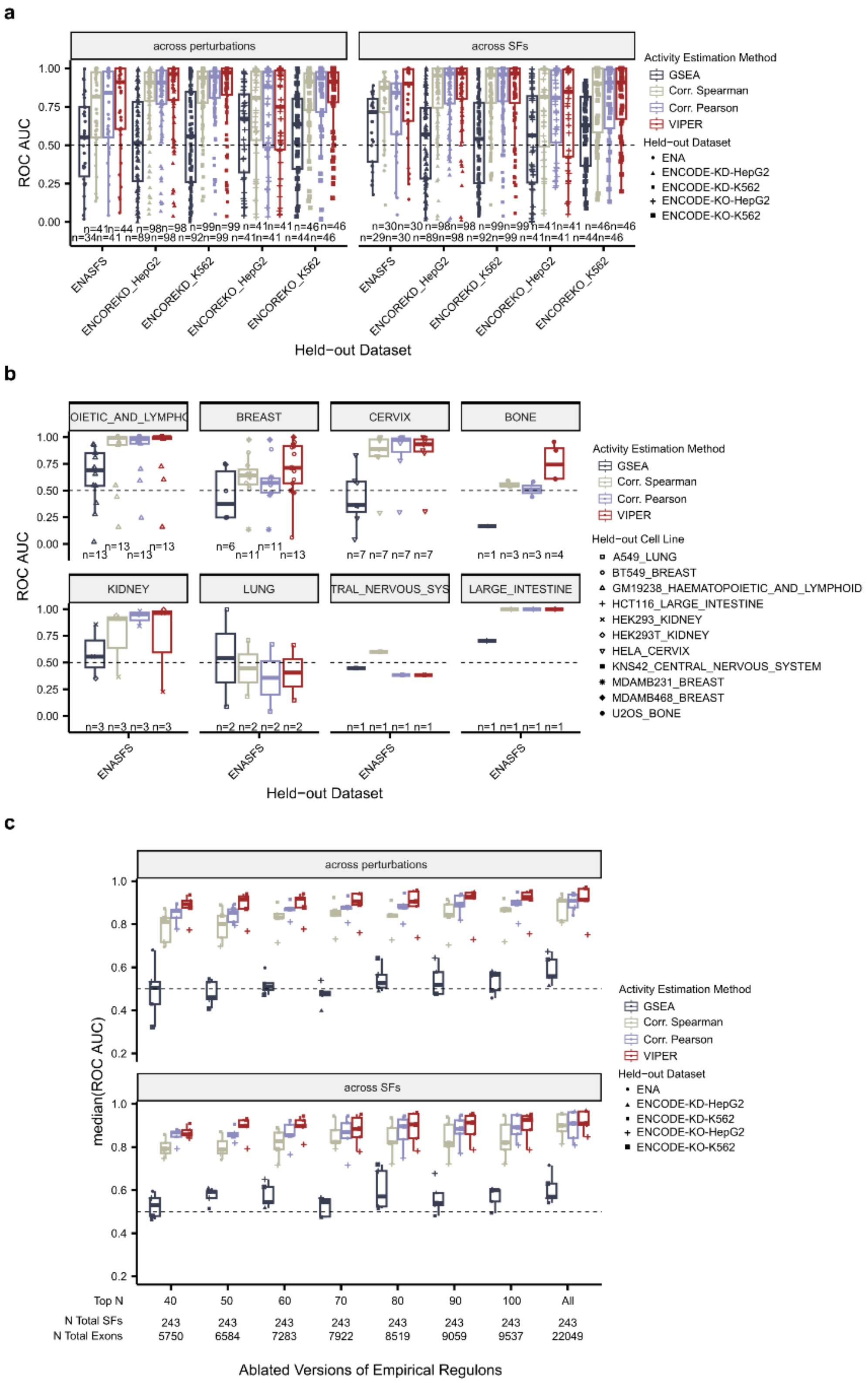

#### Supplementary Figure 2d-f

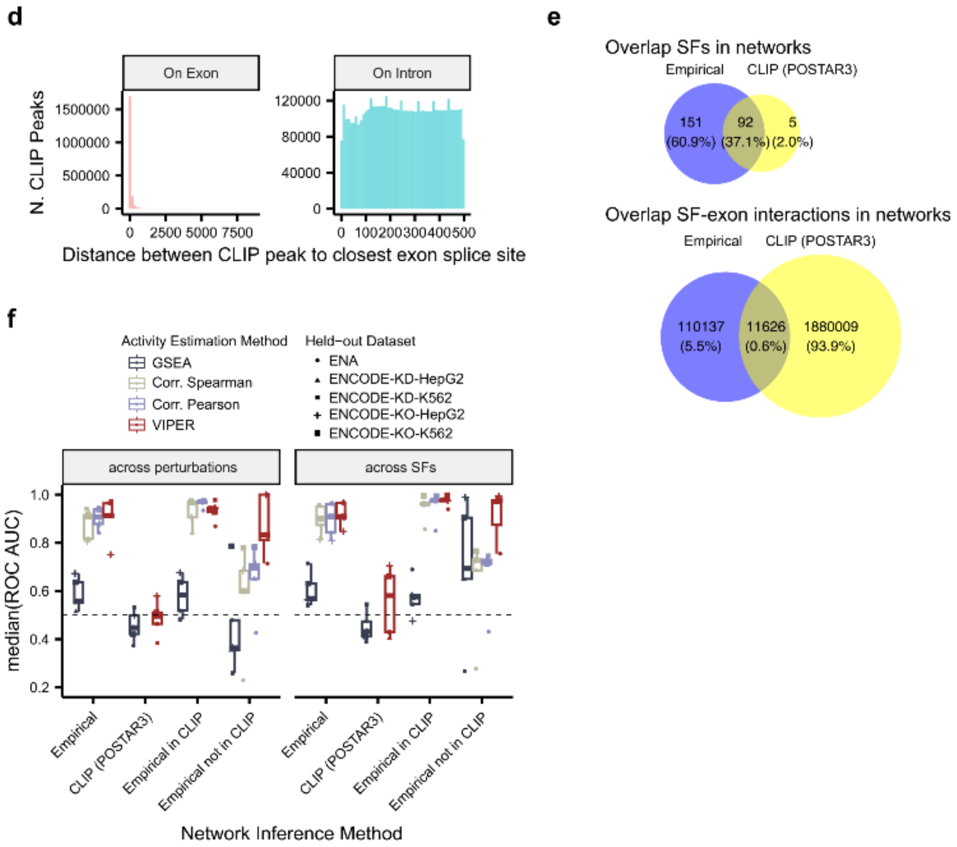


#### Supplementary Figure 3

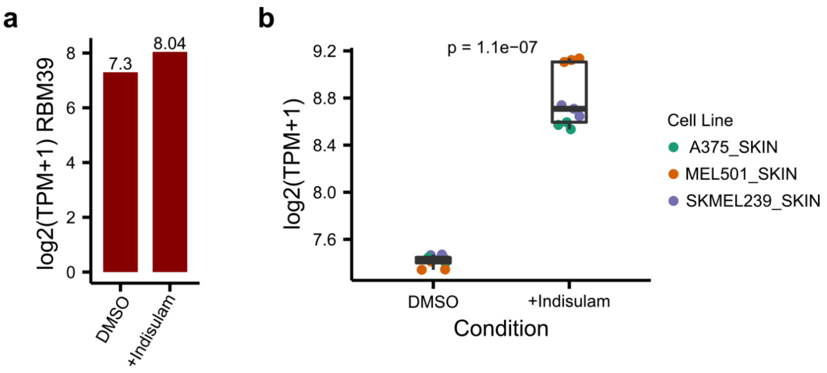

#### Supplementary Figure 4

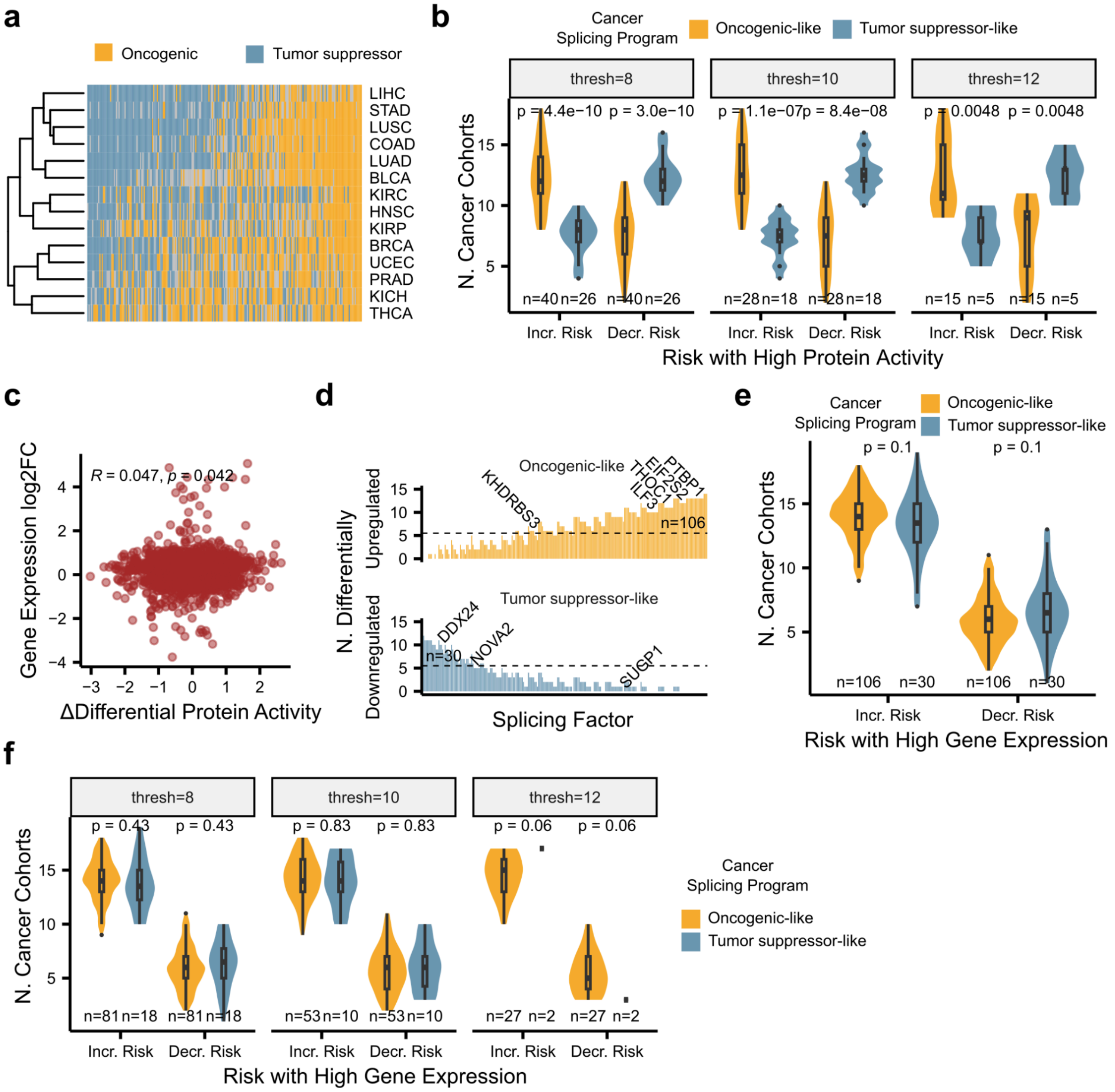

#### Supplementary Figure 5

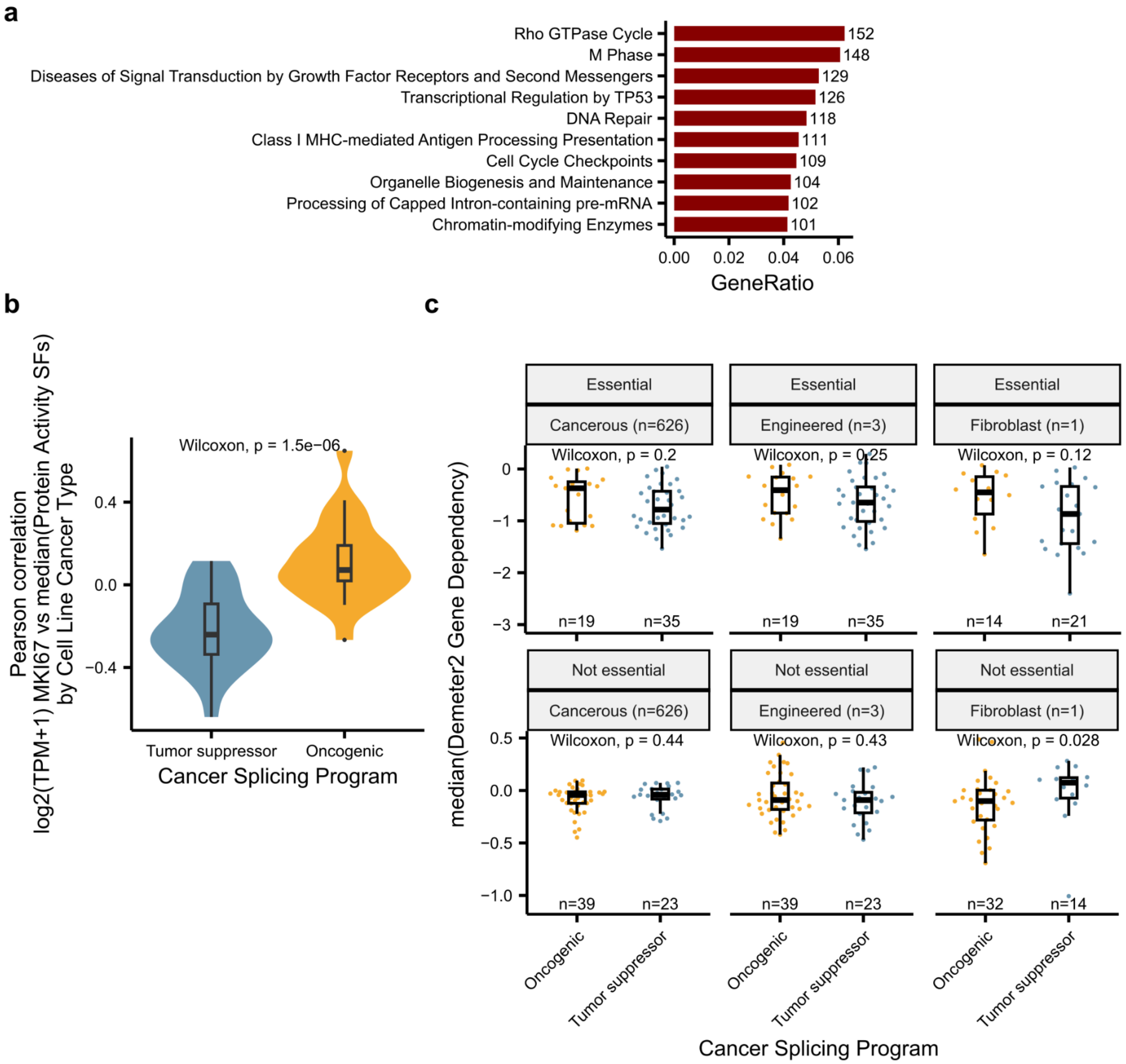

### Supplementary Figure 6

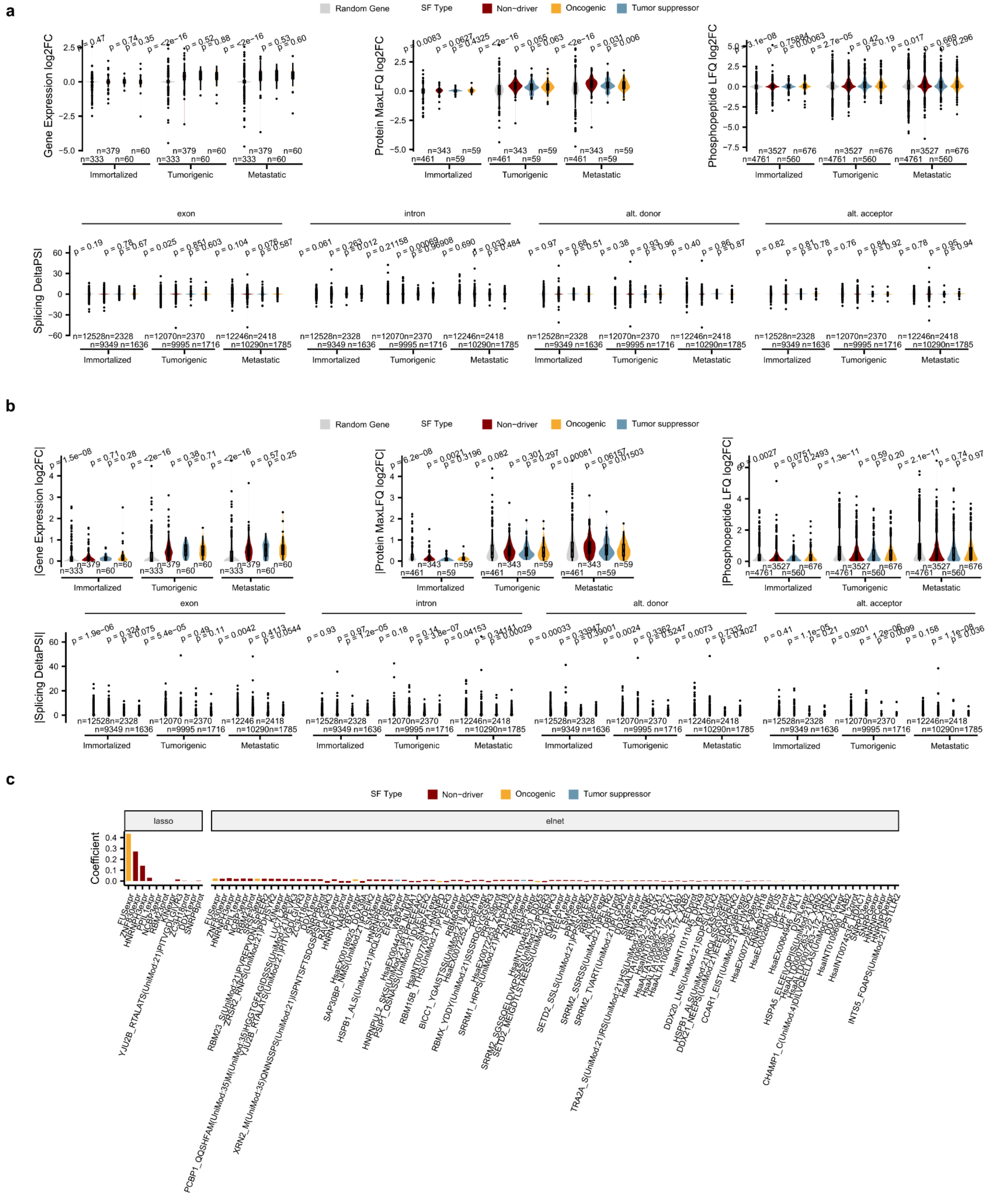

#### Supplementary Figure 7

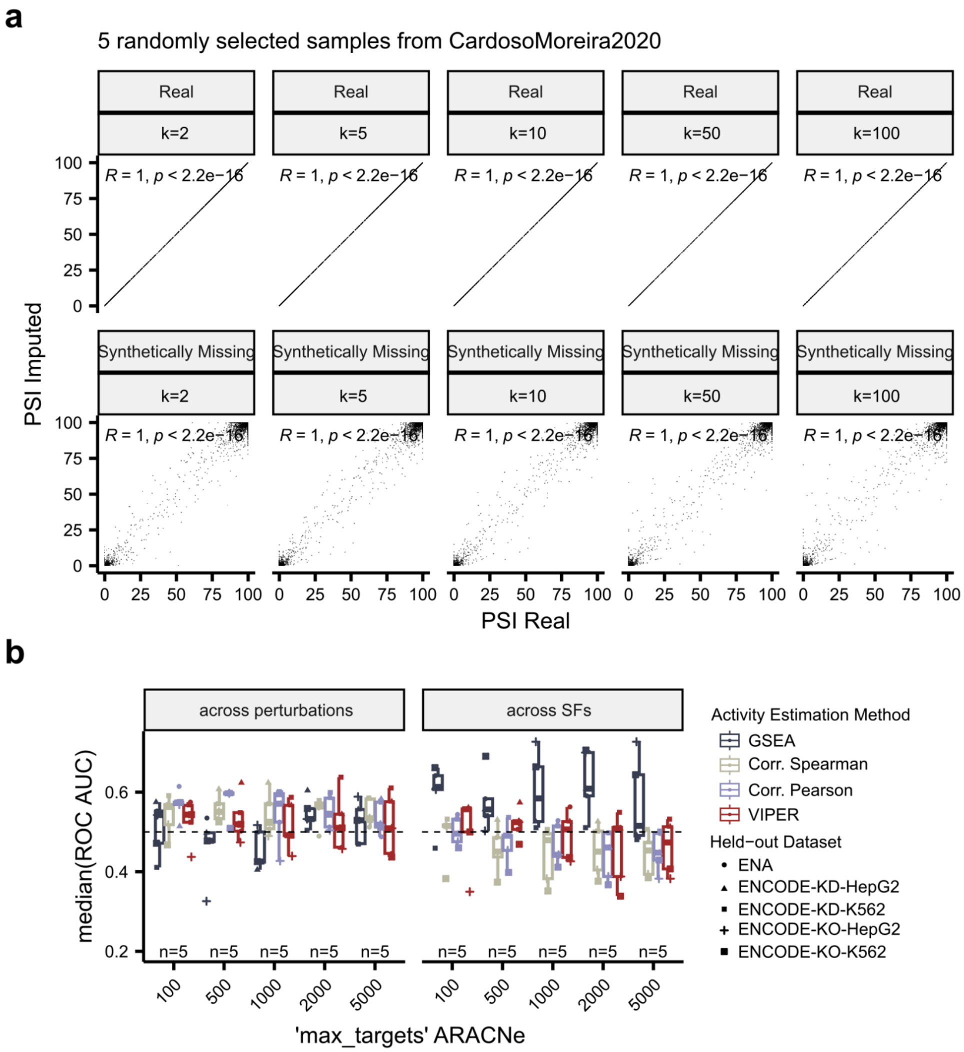

#### SUPPLEMENTARY TABLES

**Supplementary Table 1. Consensus list of 509 splicing factors.**

**Supplementary Table 2. Project identifiers of hand-curated datasets perturbing splicing factors available in ENA.**

**Supplementary Table 3. Cancer splicing programs.**

**Supplementary Table 4. Overrepresentation analysis results for the genes corresponding to target exons of each defined cancer splicing program.**
